# *C. elegans* RAP-1 reinforces LET-60/Ras induction of cell fate

**DOI:** 10.1101/297812

**Authors:** Neal R. Rasmussen, Daniel J. Dickinson, David J. Reiner

**Affiliations:** Center for Translational Cancer Research, Institute of Biosciences and Technology, Texas A&M Health Science Center, Houston, TX, 77030, USA; Department of Molecular Biosciences, University of Texas, Austin, TX, 78705; Department of Medical Physiology, College of Medicine, Texas A&M University, College Station, TX 77843, USA.

**Author notes:** Correspondence: Center for Translational Cancer Research, Institute of Biosciences and Technology, Texas A&M Health Science Center, 2121 W. Holcombe Blvd., Houston, TX 77030.

**Keywords:** Ras, Rap1, Raf, PDZ-GEF, CRISPR

## Abstract

The notoriety of the small GTPase Ras as the most mutated oncoprotein has led to a well-characterized signaling network largely conserved across metazoans. Yet the role of its close relative Rap1 (Ras Proximal), which shares 100% identity between their core effector binding sequences, remains unclear. A long-standing controversy in the field is whether Rap1 also functions to activate the canonical Ras effector, the S/T kinase Raf. We used the developmentally simpler *Caenorhabditis elegans*, which lacks the extensive paralog redundancy of vertebrates, to examine the role of RAP-1 in two distinct LET-60/Ras-dependent cell fate patterning events: induction of 1˚ vulval precursor cell (VPC) fate and of the excretory duct cell. Fluorescently tagged endogenous RAP-1 is localized to plasma membranes and is expressed ubiquitously, with even expression levels across the VPCs. RAP-1 and its activating GEF PXF-1 function cell autonomously and are necessary for maximal induction of 1˚ VPCs. Critically, mutationally activated endogenous RAP-1 is sufficient both to induce ectopic 1˚s and duplicate excretory duct cells. Like endogenous RAP-1, before induction GFP expression from the *pxf-1* promoter is uniform across VPCs. However, unlike endogenous RAP-1, after induction GFP expression is increased in presumptive 1˚s and decreased in presumptive 2˚s. We conclude that RAP-1 is a positive regulator that promotes Ras-dependent inductive fate decisions. We hypothesize that PXF-1 activation of RAP-1 serves as a minor parallel input into the major LET-60/Ras signal through LIN-45/Raf.

## Introduction

Ras, the founder of the Ras superfamily of small GTPases, is the most mutated oncoprotein in cancer (COSMIC v84; http://cancer.sanger.ac.uk/cosmic; (Papke and Der 2017). With diverse functions throughout cell biology, members of the Ras superfamily are conserved across Metazoa. Within given subfamilies, the core effector binding sequences are typically identical among *C. elegans*, *Drosophila*, and mammals, suggesting functional conservation of GTPase interactions with downstream effectors (Reiner and Lundquist 2016), many of which are also conserved across Metazoa.

Small GTPases are typically membrane-bound through lipid prenylation and processing of their C-termini (Hancock et al. 1990; Prior and Hancock 2012; Reiner and Lundquist 2016). They function as molecular switches, cycling between GTP-bound (active) and GDP-bound (inactive) states. Thus, activity of small GTPases is controlled by GEFs (<u>g</u>uanine nucleotide <u>e</u>xchange <u>f</u>actors), which displace GDP to allow GTP loading, and GAPs (<u>G</u>TPase <u>a</u>ctivating <u>p</u>roteins), which stimulate the otherwise inefficient intrinsic GTPase activity that hydrolyzes GTP to GDP (Wennerberg et al. 2005).

The Rap (<u>Ra</u>s<u>p</u>roximal) subfamily comprises GTPases closely related to Ras itself. Rap1 (previously known as K-Rev1) shares identical core effector binding sequences with Ras, while Rap2 diverges at one residue in the core effector binding sequence (Fig S1, 2). The Rap subfamily shares some GEFs and GAPs with the Ras subfamily, but also has some GEFs and GAPs that are specific to Raps (Raaijmakers and Bos 2009; Gloerich and Bos 2011). Historically, Rap1 has mostly been implicated in the regulation of cell-cell junctions, which is generally not considered to be a functional site of Ras action (Caron et al. 2000; Reedquist et al. 2000; Knox and Brown 2002; Bos 2005). Yet because of their identical core effector binding sequences, it has long been hypothesized that Ras and Rap1 share effectors and may interact in signaling networks, while this has not been suggested for Rap2. Unfortunately, an early experimental artifact in cell culture experiments clouded the role of Rap1 relative to Ras and its interaction with the canonical Ras effector, the Raf Ser/Thr kinase. Activated Rap1 transfected into cells inhibited oncogenic transformation and blocked activation of the Ras-Raf-MEK-ERK canonical MAP kinase cascade (Kitayama et al. 1989; Kitayama et al. 1990; Sakoda et al. 1992; Cook et al. 1993), yet failed to interfere with ERK activation in more physiologically relevant conditions (Zwartkruis et al. 1998). This observation led to the probably erroneous model that Rap1 was a competitive inhibitor of Ras. Because this observation was only observed under conditions of high ectopic expression (Kitayama et al. 1989), a plausible explanation is that over-expressed activated Rap1 sequestered Raf to cell-cell junctions where Ras-Raf signaling is typically not functional, thus diminishing activation of Raf-MEK-ERK signal by endogenous Ras. Yet the role of Rap1 relative to Ras-Raf signaling has remained murky since, despite certain conditions in which Rap1 appears to contribute to Raf activation in cultured mammalian cells (York et al. 1998).

However, several lines of evidence support Rap1 contributing to tumorigenic growth through activation of Raf, suggesting that Ras and Rap1 may function in parallel to activate Raf. Rap1 can oncogenically transform mammalian tissue culture cells (Altschuler and Ribeiro-Neto 1998). Non-canonical putative activating mutations in Rap1 have also been associated with Kabuki Syndrome (Bogershausen et al. 2015), part of the RASopathy spectrum of heritable disorders associated with inappropriate weak activation of the Ras-Raf signaling axis (Aoki et al. 2016). Though typically not mutated itself as an oncogene, rare mutations in Rap1 have been observed in various cancers (COSMIC v84; http://cancer.sanger.ac.uk/cosmic; Gyan et al. 2005). The paucity of such activating mutations may be because of the strong role of Rap1 in assembly and maintenance of cell-cell junctions, which may counter tumorigenic growth if not spatially controlled. A comparison to Rap1 may be drawn with the Rho GTPase family member, Rac. Oncogenic Rac mutations are rare and are mostly found in melanoma (Sergent 1990; Krauthammer et al. 2012). Yet while mostly not mutated to drive cancer, probably because of its central role in control of the cytoskeleton, cell morphogenesis, and migration, inappropriate upstream activation of Rac can still contribute to tumorigenesis (Lindsay et al. 2011; Srijakotre et al. 2017). This loss of regulation versus mutational activation has also been observed with Rap1, with the loss of a number of RapGAPs, implicating them as tumor suppressors (McLaughlin et al. 2013; Maertens and Cichowski 2014; Zhao et al. 2015).

Studies in *Drosophila* suggest that Rap1 is necessary for maximal induction of the Raf-MEK-ERK MAP kinase cascade, perhaps in parallel to Ras activation (Mishra et al. 2005; Mavromatakis and Tomlinson 2012). Yet in Drosophila, too, results with Rap1 have been contradictory (Baril et al. 2014). The *Drosophila* Roughened mutation, originally thought to be a gain-of-function mutation, was subsequently inferred to be a dominant-negative mutation (Hariharan et al. 1991; Mavromatakis and Tomlinson 2012). That *Drosophila* Rap1 is an essential gene further complicates investigation of its *in vivo* function with regard to activation of Raf during development. Consequently, there is great benefit to studying the role of Rap1 in a simple developmental system where Rap1 is not required for viability, and where there is less concern about genetic redundancy. Thus, we are investigating the role of Rap1 signaling in *C. elegans*, where Rap1 loss-of-function mutants are viable, and there is no redundancy in gene subfamilies compared to mammals (mammals possess three Ras-, two Rap1-, three Rap2-, and three Raf-encoding genes).

*C. elegans* encodes RAP-1 and RAP-2, which correspond to mammalian Rap1 and Rap2, respectively (*C. elegans* RAP-3, which has a non-conservative change in a critical residue in the core effector binding sequence, is likely to be a nematode-specific, functionally divergent Rap; Fig. S1; Reiner and Lundquist 2016). Consistent with their roles in promoting cell-cell junctions in *Drosophila* and mammalian cells, *C. elegans* RAP-1 and RAP-2 are redundant for larval molting and epithelial integrity, the double mutant causing lethality, a phenotype that is echoed by the deletion of the RapGEF, PXF-1 (Pellis-van Berkel et al. 2005). For assembly of cadherin-based cell-cell junctions in *C. elegans*, RAP-1 functions redundantly with the Ras family small GTPase, RAL-1 (<u>Ra</u>s like; Frische et al. 2007). Yet thus far there has been no examination of the role of RAP-1 in developmental patterning of the *C. elegans* vulval precursor cell (VPC) fates, a system where the sole *C. elegans* Ras ortholog, LET-60, plays a central role. Therefore, we investigated the role of RAP-1 in VPC fate patterning.

The six equipotent VPCs (P3-8.p) are induced by EGF secreted by the anchor cell (AC) in the somatic gonad, such that the VPC nearest the AC (typically P6.p) is induced to assume 1˚ fate (Fig. 1; Sternberg 2005). LET-23/EGFR receives the EGF signal, and the 1˚-promoting signal is transduced by a canonical Ras-Raf-MEK-ERK (*C. elegans* LET-60-LIN-45-MEK-2-MPK-1) MAP kinase cascade: this signal is necessary and sufficient for induction of 1˚ fate (Sundaram 2013). Induced presumptive 1˚ cells express DSL ligands (Chen and Greenwald 2004), which signal neighboring VPCs via the LIN-12/Notch receptor to assume 2˚ fate. LIN-12/Notch is necessary and sufficient for 2˚ fate (Greenwald et al. 1983; Greenwald and Kovall 2013).

**Figure 1.**
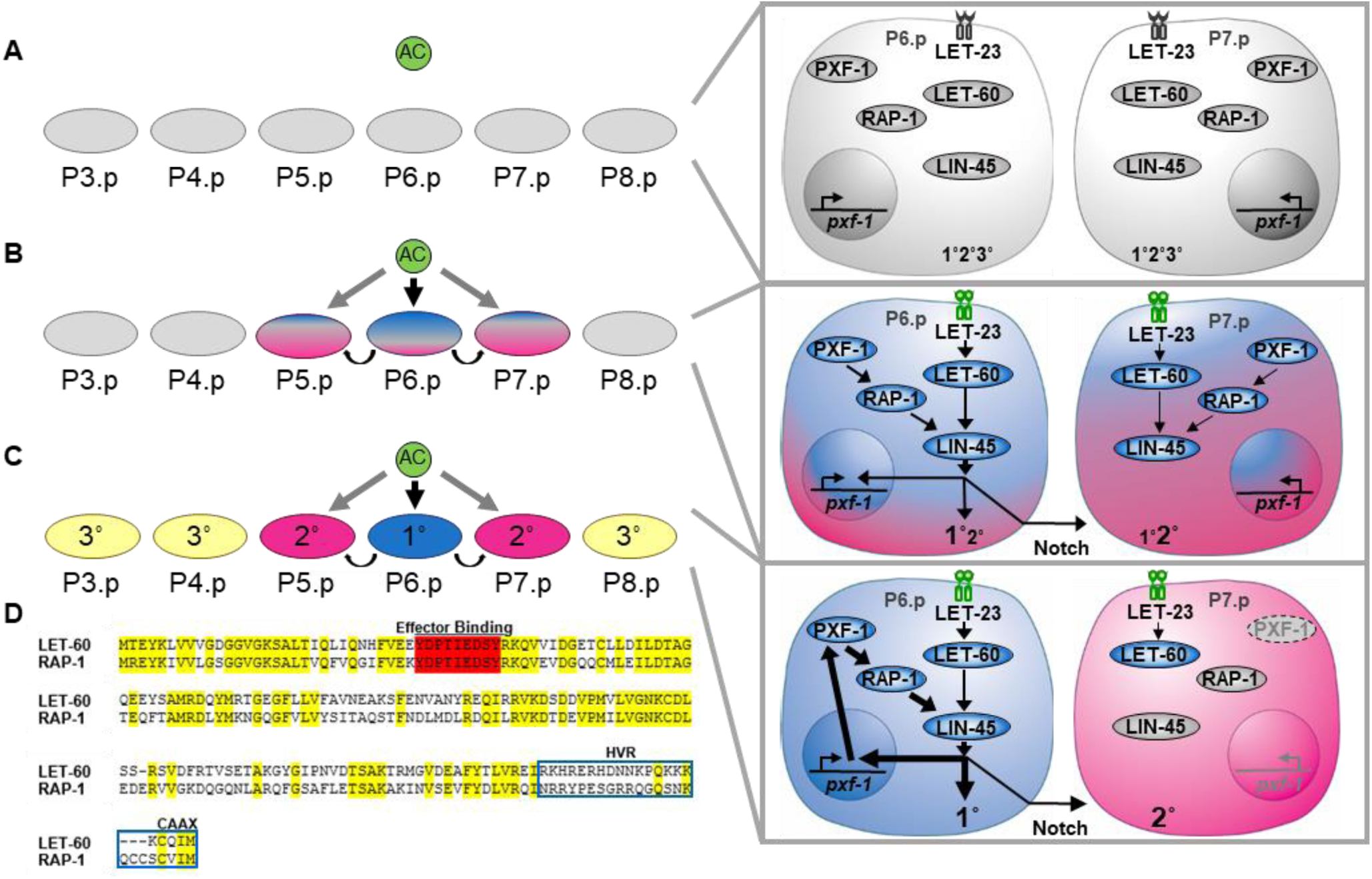
A model of RAP-1 and vulval cell fate patterning. **A)**Patterning of the vulva begins with six equipotent vulval precursor cells (VPCs). **B)** Initial cell fate specification of the VPCs begins in response to the release of EGF from the nearby anchor cell (AC) through induction of the classic LET-60/Ras-LIN-45/Raf cascade and a subsequent lateral inhibitory signal via Notch. **C)** 1˚ cell fate is then reinforced through the restricted and amplified expression of PXF-1, resulting in elevated RAP-1 activation. **D)** Sequence alignment of LET-60 and RAP-1 with the core effector binding sequence shaded in red and identical residues in yellow. Core effector binding sequences are thought to confer specificity for suites of effectors, and are generally identical across metazoans. The C-terminal hyper-variable and CAAX are outlined in the blue.

Development of the wild-type 3˚-3˚-2˚-1˚-2˚-3˚ pattern of VPC fates occurs with 99.8% accuracy (Braendle and Felix 2008). During VPC fate patterning, cells are initially specified, then become committed to their fate (Sternberg 2005). During this process, the VPC signaling network is at least partially re-programmed, perhaps contributing to final commitment and fidelity. For example, after initial induction, LIN-12/Notch is internalized and degraded in presumptive 1˚ cells (Shaye and Greenwald 2002, 2005), thus precluding 2˚-promoting signal in a cell that is specified to 1˚ fate. Conversely, presumptive 2˚ cells express the LIN-12/Notch transcriptional client, LIP-1/ERK phosphatase (Berset et al. 2001; Yoo et al. 2004), thereby prohibiting 1˚-promoting signaling in cells that are specified as 2˚. Additionally, expression of a suite of other LIN-12/Notch transcriptional client genes is altered after initial induction (Berset et al. 2001; Yoo et al. 2004; Berset et al. 2005; Yoo and Greenwald 2005; Zhang and Greenwald 2011), supporting the idea of network reprogramming. We observed further evidence consistent with VPC re-programming with *ral-1* promoter expression – initially expressed in all VPCs – subsequently being excluded from presumptive 1˚ cells, thereby prohibiting 2˚-promoting signaling in from cells that are specified as 1˚ (Reiner 2011; Zand et al. 2011). Most of this re-programming at the transcriptional level occurs prior to the first VPC division. Though we do not know the precise point at which terminal commitment occurs (Wang and Sternberg 1999), we hypothesize that transcriptional and potentially post-translational network re-programming, occurring prior to the first cell division, is a critical component of the high fidelity observed in this developmental decision.

In this study, we use VPC fate patterning to investigate the long-standing question of the relationship between Ras and Rap1. We used CRISPR to fluorescently tag the endogenous RAP-1 and found it to be expressed throughout the animal and localized to plasma membranes and with increased concentration at cell-cell junctions. We find that deletion of RAP-1 in an otherwise wild-type background result in mild and low penetrance VPC patterning defects. Use of sensitized genetic backgrounds indicated a cell-autonomous role for RAP-1 in promoting 1˚ fate. CRISPR-mediated mutational activation of endogenous RAP-1 revealed that RAP-1 is sufficient to induce ectopic 1˚ cells, and also induce duplication of the excretory duct cell, another cell induction event dependent on LET-60/Ras-LIN-45/Raf signaling. We found that the PXF-1/PDZ-GEF is also required for maximal 1˚ induction, consistent with a role for PXF-1 as the activating RAP-1 GEF in VPC patterning. Furthermore, we find that during response to EGF signal, transgenic GFP expression from the *pxf-1* promoter changes from uniform expression across VPCs to increased in presumptive 1˚ cells while decreased in presumptive 2˚ cells. This observation is another confirmation of the re-programming hypothesis and suggests that the activation of RAP-1 is restricted to 1˚ cells during the phase when initial patterning of VPCs is reinforced. Taken together, our results support a model in which a PXF-1-RAP-1 signal functions as a minor parallel input into the major LET-60-LIN-45 1˚-promoting signaling.

## Materials and Methods

### C. elegans handling and genetics

All strains were derived from the parent N2 wild-type strain. Animals were grown at 20˚C under standard culturing conditions on NGM agar plates seeded with OP50 bacteria, unless stated otherwise (Brenner 1974). Crosses were performed using standard methods. Strain details are shown in Supplementary Table 1.

The *let-60(n1046*gf) strain undergoes genetic drift upon continuous growth, resulting in increased phenotype strength (Zand et al. 2011). We therefore established firm guidelines to ensure consistent results. Immediately upon thawing or construction, strains harboring *n1046* were scored and then starved and parafilmed as a reference strain. For subsequent experiments, we reestablished growing strains each week to avoid drift while in continuous culture. Experiments were only reported if the *let-60(n1046*gf) control was within the well-validated baseline levels of induction. We did not observe any similar drift in *rap-1(re180*gf) or *let-23(sa62*gf) strains. Strains harboring homozygous *let-23(sa62*gf) mutations did exhibit delayed growth, necessitating scoring VPC induction a day later than with other strains.

### Plasmids, Generation of CRISPR strains

Details of plasmid constructions are available upon request. Plasmids used are shown in Supplementary Table 4. sgRNA sequences and repair templates are listed in Supplementary Tables 5 and 6, respectively. The *rap-1(re180*gf) CRISPR strain was generated using the co-CRISPR strategy (Paix et al. 2014) by microinjection of pJA58 (50 ng/ul), pNR21 (50 ng/ul), *rap-1* (G12V) ssODN repair template (500 uM), *dpy-10(cn64)*, ssODN repair template (500 uM)) and co-injection marker pPD118.33 (20 ng/ul) into N2 wild-type animals. Primers and conditions for PCR genotyping are listed in Supplementary Table 3. PCR detection of *rap-1(re180*gf) was determined using an overnight digestion with BamHI (NEB), NEB Cutsmart buffer, and water to a final volume of 50 ul.

### Scoring vulval induction and fate reporters (VPC and excretory duct)

To score the formation of normal and ectopic vulval induction, late L4 animals were mounted in M9 on slides with a 3% NG agar pad containing 5mM sodium azide and examined by DIC/Nomarski optics (Nikon eclipse Ni). To ensure accuracy and reproducibility, all presented data are from animals grown together and are representative non-pooled samples. Presence of the 1˚ VPC transcriptional reporters *arIs92*[P_*egl-17::*_*CFP-LacZ*] and *arIs131*[P_*lag-2::*_*2xNLS::YFP*] were scored for each of the six VPCs with the Nikon Eclipse Ni microscope, with a Nikon DS-Fi2 color camera and using the NIS Elements Advanced Research, Version 4.40 (Nikon) software package. L3 animals were scored at the Pn.px (two-cell) stage to ensure that animals were past the point of EGF induction of VPCs. Using the same equipment, newly hatched L1 larvae were scored for the presence of a single or duplicated excretory duct cell(s) using the *saIs14*[P_*lin-48*_::GFP] transgene (Johnson et al. 2001).

### RNA interference (RNAi)

Bacterially mediated RNAi experiments were conducted at 23˚C on NGM agar plates supplemented with 1 mM IPTG and 50 µg/ml carbenicillin as described (Kamath and Ahringer 2003). In our hands, we typically obtain more robust RNAi results at 23˚C than at 20˚C (Zand et al. 2011). All RNAi clones were sequenced to confirm identity. RNAi plates were seeded with 80 µl HT115 bacteria harboring clones of *C. elegans* genes (Source Bioscience) or the negative control *luciferase* (*luc*; Shin *et al*., in preparation) and allowed to grow overnight at room temperature. Late L4 larvae were added to each plate and transferred to a fresh RNAi plate the next day. Founding parents were then removed the following day. Late L4 progeny were scored for the formation of the principal vulva and/or ectopic pseudovulvae by DIC/Normarski optics 2 days later. However, as noted above, strains homozygous for the *let-23(sa62*gf) allele reach maturity at a slower rate, and consequently were scored 4 days after parents were removed. For each experiment, *pop-1(RNAi)*, which confers 100% embryonic or L1 lethality under optimal conditions, was used as a control for maximal RNAi efficacy; animals were only scored from experiments in which 100% lethality was observed. RNAi strains used are included in Supplemental Table 2.

VPC-specific RNAi was performed using the genotype *let-23(sa62*gf*); rde-1(ne219); mfIs70[P*_*lin-31*_::*rde-1(+)*, P_*myo-2*_*::gfp]*. The *mfIs70[P*_*lin-31*_::*rde-1(+), P*_*myo-2*_::*gfp]* integrated transgene which rescues the loss of *rde-1* specifically in the VPCs (Barkoulas et al. 2013) was introduced and tracked in crosses using the *dpy-11(e224) unc-76(e911)* chromosome in *trans* as a balancer.

### Fluorescent microscopy (imaging)

Animals were mounted live in M9 buffer on slides with a 3% agar pad containing 5 mM sodium azide. DIC/Nomarski optics and epifluorescence microscopy were captured using the Nikon Eclipse Ni and confocal images using the A1si Confocal Laser Microscope (Nikon). Captured images were processed using NIS Elements Advanced Research, Version 4.40 (Nikon) and Adobe Photoshop CC 2018 software packages (inverted images).

Plasmids and strains are available upon request. The authors affirm that all data necessary for confirming the conclusions of the article are present within the article, figures, and tables.

## Results

### RAP-1 is expressed in VPCs and localized to plasma membrane and junctions

The Ras superfamily of small GTPases is characterized by lipid modification of their C-terminal CAAX motif, resulting in their subcellular localization to the lipid bilayers. For Ras itself, this primarily means localization to the plasma membrane (Hancock et al. 1990; Prior and Hancock 2012; Reiner and Lundquist 2016). Likewise, Rap1 has been observed to be found on the plasma membrane (Pizon et al. 1994). Prior work reported LET-60/Ras to ubiquitously expressed in *C. elegans* (Dent and Han 1998). Although the *let-60/Ras* promoter fusion could not provide information regarding subcellular localization, ectopic induction of 1˚s conferred by the *let-60(n1046*gf) activating mutation was reversed by treatment of farnesyl transferase inhibitors, which block C-terminal prenylation and hence activity (Hara and Han 1995). Consequently, we are reasonably confident that *C. elegans* LET-60 is targeted to the plasma membrane.

To determine the expression pattern and subcellular localization of endogenous RAP-1, we used CRISPR-Cas9-mediated genome editing to insert an mNeonGreen::3xFlag epitope tag into the 5’ end of the endogenous *rap-1* gene, generating *rap-1(cp151[mNeonGreen^3xFlag::rap-1])* (Fig. S3; Dickinson et al. 2015). In keeping with previous observations with *let-60*, tagged RAP-1 expression was ubiquitous throughout the animal during development, and localized to the plasma membrane (Fig. S4). Of particular importance to this study, RAP-1 expression was observed in the VPCs prior to EGF induction (1 cell stage, Pn.p; Fig. 2A,C), after EGF induction (2 cell stage, Pn.px; Fig. 2B,D), and throughout vulval morphogenesis.

**Figure 2.**
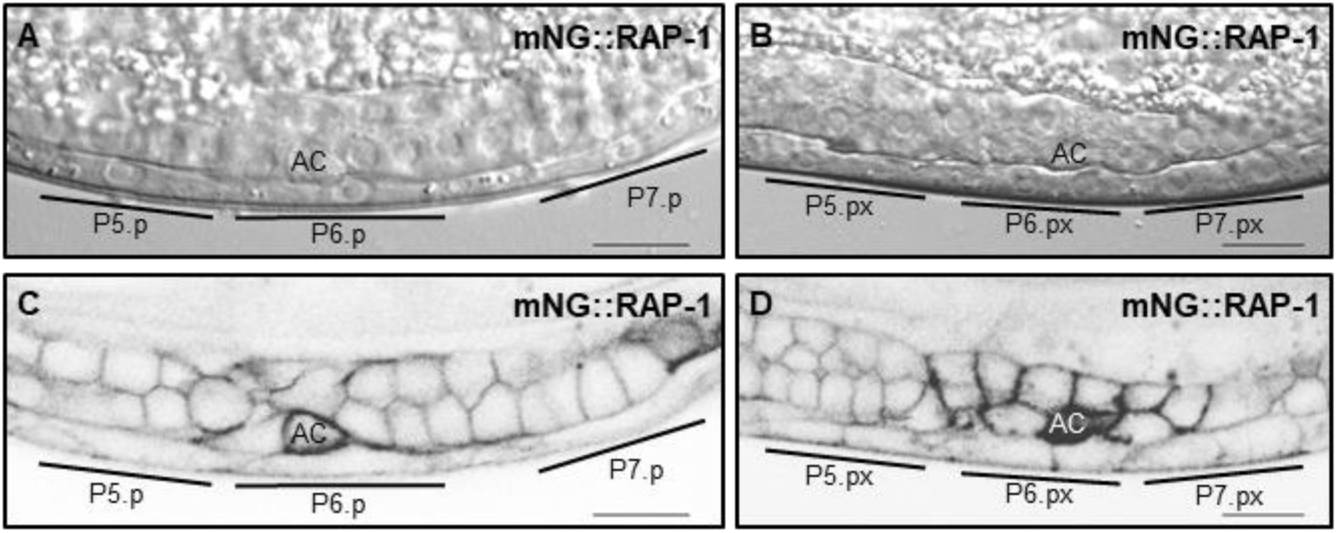
Endogenous RAP-1 is expressed throughout the VPCs. Representative DIC **(A, B)** and confocal fluorescence **(C, D)** micrographs of *rap-1(cp151[mNG^3xFlag::rap-1])* at **(A, C)** the 1-cell (Pn.p) and **(B, D)** 2-cell (Pn.px) stages.”mNG” = mNeonGreen (Shaner, Lambert et al. 2013). Scale bar = 10 µm.

We also observed that tagged endogenous RAP-1 is localized to cell-cell junctions between hypodermal seam cells (Fig. S4A). This observation is consistent with the established role of RAP-1 in the assembly of cadherin complexes at junctions during embryogenesis, a process that is redundant with another small GTPase, RAL-1 (Frische et al. 2007), and our observation that tagged endogenous RAL-1 also localizes to the plasma membrane and cell-cell junctions (Shin *et al*., submitted). This finding also conforms with the established role of Rap1 in junctional biology in *Drosophila* and mammalian cell culture (Caron et al. 2000; Reedquist et al. 2000; Knox and Brown 2002).

Intriguingly, we observed increased expression of tagged endogenous RAP-1 in the anchor cell (AC). The AC is known to undergo polarized invasive behavior, directed toward the P6.p/1˚ lineage of the vulva (Sherwood and Sternberg 2003; Hagedorn et al. 2009; Ziel et al. 2009) Fig. 2C,D). We additionally observed increased expression of RAP-1 in the distal tip cell (DTC) of the developing somatic gonad (Fig. S4B), a tissue that also goes through invasive migration. However, our later experiments indicate that RAP-1 functions cell autonomously to regulate VPC fate patterning, so we interpret the increased RAP-1 expression in the AC to be unrelated to VPC fate patterning.

### RAP-1 is necessary for maximal induction of 1˚ VPCs

The early false lead in mammalian cell culture of Rap1 as a competitive inhibitor of Ras has long muddied the waters of our understanding of the role of Rap1 in Ras signaling (Kitayama et al. 1989; Kitayama et al. 1990; Sakoda et al. 1992; Cook et al. 1993; see Introduction). Additionally, investigations of Rap1 have been complicated by its multiple isoforms in vertebrates and its essential role in development in *Drosophila* (Mishra et al. 2005). Previous work showed *C. elegans rap-1* putative null mutant animals to be fertile and viable, including *mNeonGreen^SEC^3xFlag::rap-1* animals retaining the SEC positive/negative selection cassette, which should disrupt RAP-1 expression (Frische et al. 2007; Dickinson et al. 2015). We more closely examined animals mutant for two independent *rap-1* strong loss or null alleles. Both *rap-1(tm861)* (a deletion that results in a frame shift) and *rap-1(pk2082)* (nonsense allele in exon 5 of 6; Fig. S3) conferred low penetrance abnormal vulval patterning, which we interpret to be missing 2˚ cells (Fig. S5). This phenotype is similar to that observed in animals mutant for the hypomorphic *lin-3(e1417)*, *let-60(n2021)* and *lin-45(n2506)* mutations, with reduction of function of EGF, Ras and Raf orthologs, respectively (Wang and Sternberg 1999; Zand et al. 2011). Retrospectively, we interpret this low penetrance under-induction of 2˚ cells to be due to weak specification of 1˚ cells. Indeed, the mutations mentioned here, which confer phenotypes stronger than those of *rap-1* mutations, frequently confer a lack of 1˚ cells altogether. We speculate that these under-induced 1˚ cells express insufficient levels of the triply redundant DSL ligands, LAG-2, APX-1, and DSL-1 (Chen and Greenwald 2004) Zhang and Greenwald 2011) to reliably induce the neighboring VPCs to become 2˚.

The low penetrance of these *rap-1* mutant defects makes the proper interpretation of the function of RAP-1 in VPC fate patterning difficult. Hence, we used a sensitized genetic background, *let-23(sa62*gf) (Katz et al. 1996), which is sufficient to induce ectopic 1˚s and are sensitive to perturbation of modulatory signals (Zand et al. 2011). The addition of *rap-1(tm861)* resulted in significantly decreased ectopic 1˚s compared to *let-23(sa62*gf) alone (Fig. 3A). We also targeted *rap-1* loss in a VPC-specific manner using the *let-23(sa62*gf); *rde-1(ne219)* background with the *mfIs70* integrated transgene (P_*lin-31*_ driving VPC-specific rescue with *rde-1(+)*). In agreement with our previous findings, VPC-specific depletion of *rap-1* resulted in a significant decrease in the formation of ectopic 1˚s (Fig. 3B). These results demonstrated that RAP-1 functions cell-autonomously in the VPCs. To further evaluate the effects of *rap-1* loss, we utilized the *let-60(n1046gf*) mutant strain, which harbors the G13E mutation analogous to mutation in mammalian Ras that lock the protein in the GTP-bound form, thus causing the protein to be constitutively active (Han et al. 1990; Beitel et al. 1990). Similar to our findings with *let-23(sa62*gf), above, *rap-1(tm861)* reduced ectopic 1˚ induction in the *let-60(n1046*gf) background (Fig. 3C). Consequently, we hypothesize that RAP-1 functions in parallel to or downstream of LET-60 to promote 1˚ fate. Given the biochemical relationships of mammalian Rap1 and Ras to Raf, we favor an interpretation of parallelism, potentially converging on LIN-45/Raf.

**Figure 3.**
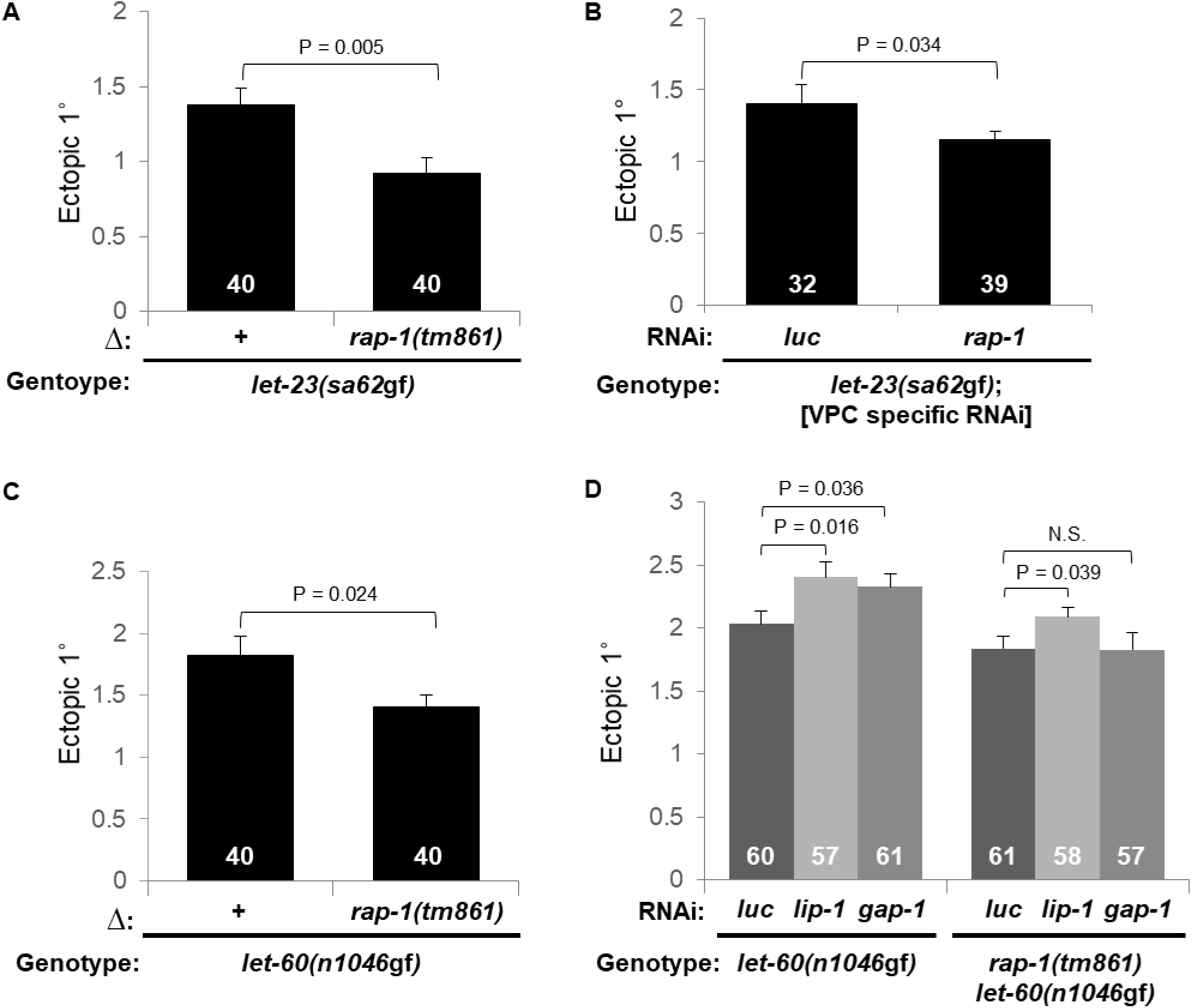
RAP-1 is necessary for maximal 1˚ induction and confers response to GAP-1/RasGAP. **A)** Deletion or **B)** VPC-specific RNAi knockdown of *rap-1* decreased ectopic 1° induction in the *let-23(sa62*gf) (*cis*-marked with *unc-4(e120)*) background. VPC-specific RNAi = *rde-1(ne219)*; *mfIs70*[P_*lin-31*_::*rde-1*(+), P_*myo-2*_::*gfp*]. Luciferase-directed (*luc*) RNAi served as a negative control (Shin *et al*., submitted). **C)** *rap-1* deletion decreased ectopic 1° induction in the *let-60(n1046*gf) background. **D)** *let-60(n1046*gf) animals are sensitive to *gap-1*- and *lip-1-*directed RNAi (depletion of RasGAP and ERK phosphatase, respectively), while *rap-1(tm861) let-60(n1046*gf) animals are insensitive to *gap-1(RNAi)* but still responsive to *lip-1(RNAi)*. All data are representative non-pooled assays, collected on the same day, of the mean ectopic 1˚ cells ± S.E.M. **D)** *let-60(n1046*gf) vs. *rap-1(tm861) let-60(n1046*gf) animals were scored on separate days. Y-axes indicate number of ectopic 1˚ cells. N = white numbers. P values were calculated by T-test or ANOVA.

We previously noted that *let-60(n1046*gf), which is predicted to be GAP insensitive, was paradoxically still responsive to RNAi-mediated depletion of the negative regulator, GAP-1 (Zand et al. 2011). We reproduced this result, observing that the *let-60(n1046*gf) single mutant responded to *gap-1(RNAi)* by increasing ectopic 1˚ induction, with RNAi targeting the LIP-1/ERK phosphatase (Zand et al. 2011) as a positive control and *luciferase*-directed RNAi as a negative control (Fig. 3D). However, we found that *rap-1(tm861) let-60(n1046*gf) double mutant animals no longer responded to *gap-1(RNAi)* while still responding to *lip-1(RNAi)* (Fig. 3D). These results suggest that GAP-1 functions to promote GTP hydrolysis of both RAP-1 and LET-60/Ras related small GTPases. Accordingly, the mammalian GAP1 subfamily of GAPs, orthologous to *C. elegans* GAP-1 (Stetak et al. 2008; Grewal et al. 2011), has been shown to be bifunctional, targeting mammalian Ras as well as Rap1 and Rap2 as substrates (Kupzig et al. 2006).

### RAP-1 is sufficient to induce ectopic 1˚ cells

If RAP-1 functions in parallel to LET-60 to activate LIN-45/Raf and hence induction of 1˚ cells, we would expect constitutively active RAP-1 to promote induction of ectopic 1˚ cells. We used CRISPR/Cas9 genome editing to introduce the classical G12V activating mutation into the endogenous *rap-1* locus (Fig. 4A). The G1 region in both Ras/Rap1 proteins is exceptionally well conserved across species (Fig. S1,2; Wennerberg et al. 2005; Reiner and Lundquist 2016) allowing the G12V mutation to be successfully used for both Ras and Rap1 in *C. elegans* (Pellis-van Berkel et al. 2005; Zand et al. 2011) and *Drosophila* (Mishra et al. 2005), as well as mammalian cell culture (Vossler et al. 1997). The *rap-1(re180*gf) allele can be detected by PCR and restriction enzyme digest due to the introduction of the concomitant silent BamHI site (Fig. 4B).

**Figure 4.**
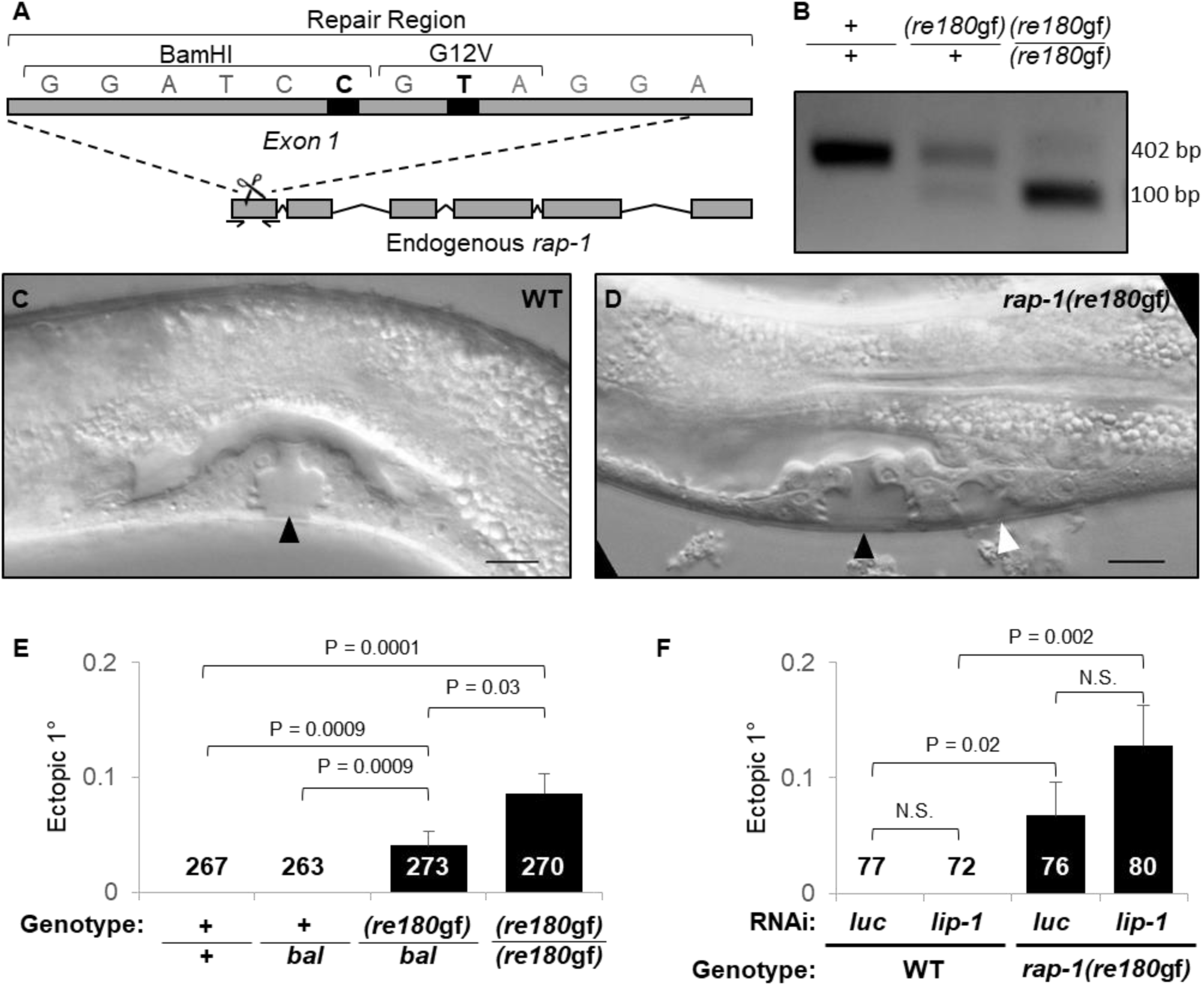
RAP-1 is sufficient to promote ectopic 1° cells. **A)** Diagram of the strategy for CRISPR/Cas9-dependent knock-in of the activating (G12V) mutation into endogenous *rap-1*, along with a silent BamHI restriction site for genotyping. **B)** PCR genotyping of the *rap-1(re180*gf) mutant allele, followed by BamHI digestion and agarose gel electrophoresis. **C,D)** Representative DIC micrographs at the late L4 stage shows the wild-type vulva (black triangle) in **(C)** wild-type (WT) animals, and induction of ectopic 1°s (white triangle) in **(D)** *rap-1(re180*gf). Scale bar = 10 µm. **E)** Both heterozygous and homozygous *rap-1(re180*gf) animals exhibited increased ectopic 1° induction compared to WT. *bal* = *rap-1* balancer *nT1qIs51[*P_*myo-2*_::*gfp*, P*pes-10::gfp*, P*F22B7.9::gfp]*. **F)** RNAi depletion of *lip-1* did not significantly increase ectopic 1°s (*luc = luciferase* RNAi negative control). Y axis = mean number of ectopic 1˚ VPCs ± S.E.M. N = numbers on columns. P values were calculated by T-test.

*rap-1(re180*gf) is sufficient to promote ectopic 1˚ cell differentiation (Fig. 4C, D, S6A). As predicted based on *let-60(n1046*gf) (Beitel et al. 1990; Han et al. 1990), the *rap-1(re180*gf) mutation acts in a gain-of-function manner, evidenced by semi-dominant induction of ectopic 1˚ cells (Fig. 4E). Depletion of the LIP-1/ERK phosphatase can increase ectopic 1˚ induction in sensitized backgrounds (Berset et al. 2001; Yoo et al. 2004; Berset et al. 2005 Zand et al. 2011), and we used *lip-1(RNAi)* as a positive control in this study (Fig. 3D). Unexpectedly, *lip-1(RNAi)* failed to enhance the ectopic 1˚ induction in the *rap-1(re180*gf) background (Fig. 4F).

To verify that *rap-1(re180*gf) was promoting 1˚ VPC transcriptional programs, we utilized two validated reporters of 1˚-promoting signaling, *arIs131 (*P_*lag-2::*_*YFP*, P_*ceh-22*_::GFP) and *arIs92 (*P_*egl-17*_::CFP-LacZ, P_*ttx-3*_::GFP) (Yoo et al. 2004; Zhang and Greenwald 2011). In control animals, expression of P_*lag-2*_*::YFP* was only observed in P6.px, compared to *rap-1(re180gf)* mutants where we also observed YFP in both P5.px and P7.px (Fig. 5A-D). A similar increase in ectopic expression of P_*egl-17*_::CFP-LacZ was observed with P_*egl-17*_*::cfp-LacZ* with the addition *rap-1(re180*gf) (Fig. S6B-E). For both expression of P_*lag-2*_::*YFP* and P_*egl-17*_::CFP-LacZ the addition of *rap-1(re180gf)* resulted in a significant increase expression outside of P6.px. Rare expression in and transformation of 3˚ VPCs was also observed (data not shown; Fig. S6A).

**Figure 5.**
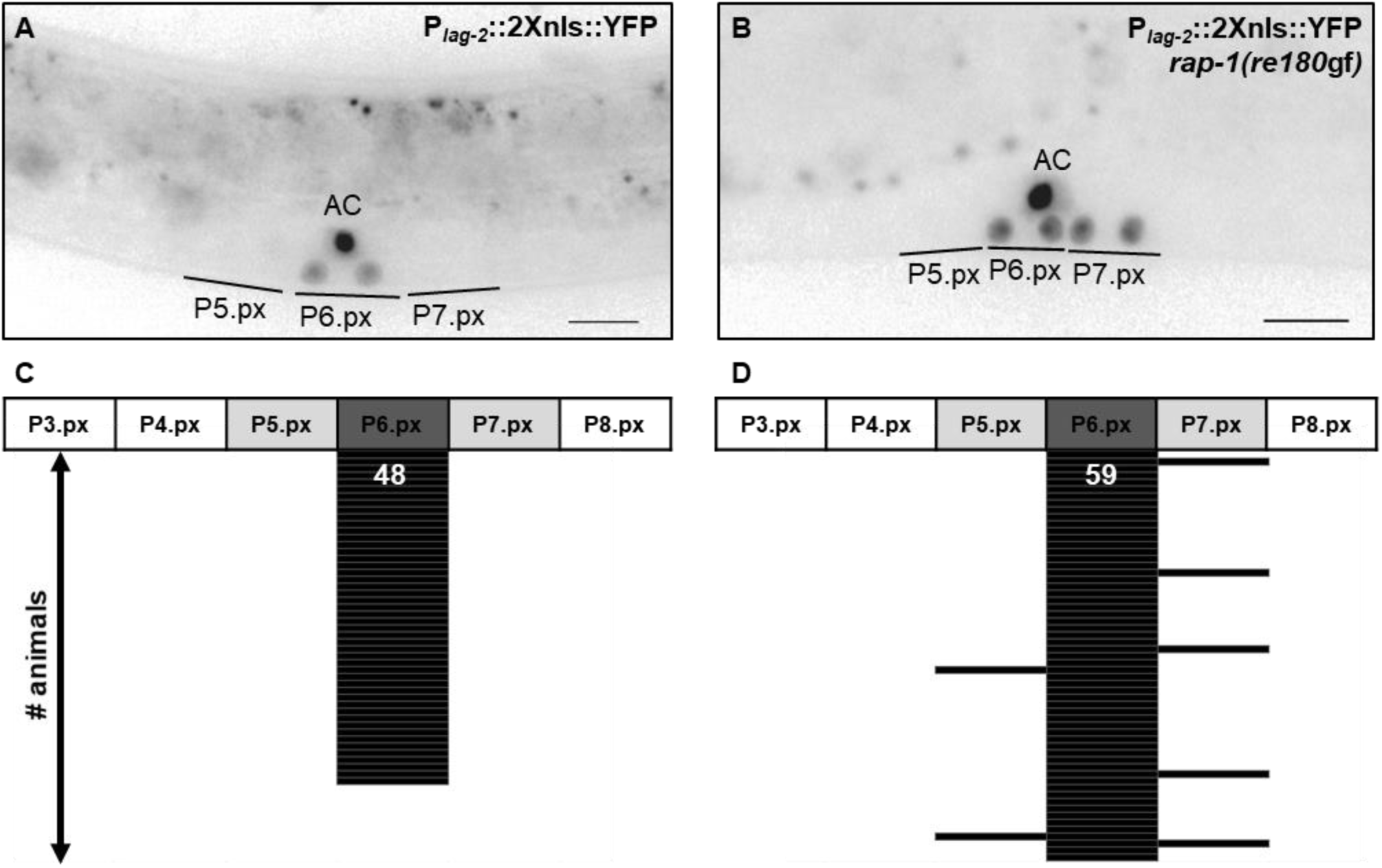
RAP-1 activation results ectopic expression of 1° VPC transcriptional reporters. **A,B)** Representative epifluorescence micrographs in animals at the 2-cell (Pn.px) stage show the inappropriate expression of 1° signaling reporter *arIs131*[P_*lag-2*_::2Xnls::YFP] in **(B)** *rap-1(re180*gf) but not **(A)** wild type. Scale bar = 10 µm. **I,J)** Schematic representation of the expression of the *arIs131*[P_*lag-2*_::2Xnls::YFP] signaling reporter across the all six VPCs for both **(C)** control and **(D)** *rap-1(re180*gf). The addition of *rap-1(re180gf)* resulted in a significant increase of expression outside of P6.px (P = 0.016) P values were calculated by Fisher’s Exact test. N = white number, with each line representing an animal.

### RAP-1 is sufficient to promote excretory duct cell duplication

We observed that some *rap-1(re180*gf) animals exhibited a ventral protrusion near the posterior bulb of the pharynx (Fig. 6B). *let-60(n1046*gf) confers a similar ventral protrusion, which was found to be duplication of the excretory duct cell (Beitel et al. 1990; Han et al. 1990; Yochem et al. 1997). Conversely, defective 1˚-inducing cascade, from LIN-3/EGF to the LET-60/Ras-LIN-45/Raf MAP kinase cascade, confers a high degree of “rod-like” larval lethality due to failure of the excretory duct cell to develop, and hence the inference of inability to excrete solutes (Beitel et al. 1990; Han and Sternberg 1990; Han et al. 1990; Yochem et al. 1997). To determine if RAP-1 activation is sufficient to result in excretory duct cell duplication, we used the *saIs14 (*P_*lin-48*_::*GFP)* transgene, a cell fate reporter for the duct cell (Johnson et al. 2001). Mirroring our results in VPC cell fate patterning, *rap-1(re180*gf) was sufficient to drive duct-cell duplication, but at a reduced rate in comparison to *let-60(n1046*gf) (Fig. 6). We observed no rod-like larval lethality in *rap-1(tm861)* or *rap-1(pk2082)* mutants, suggesting that *rap-1* is not necessary for induction of the excretory duct cell.

**Figure 6.**
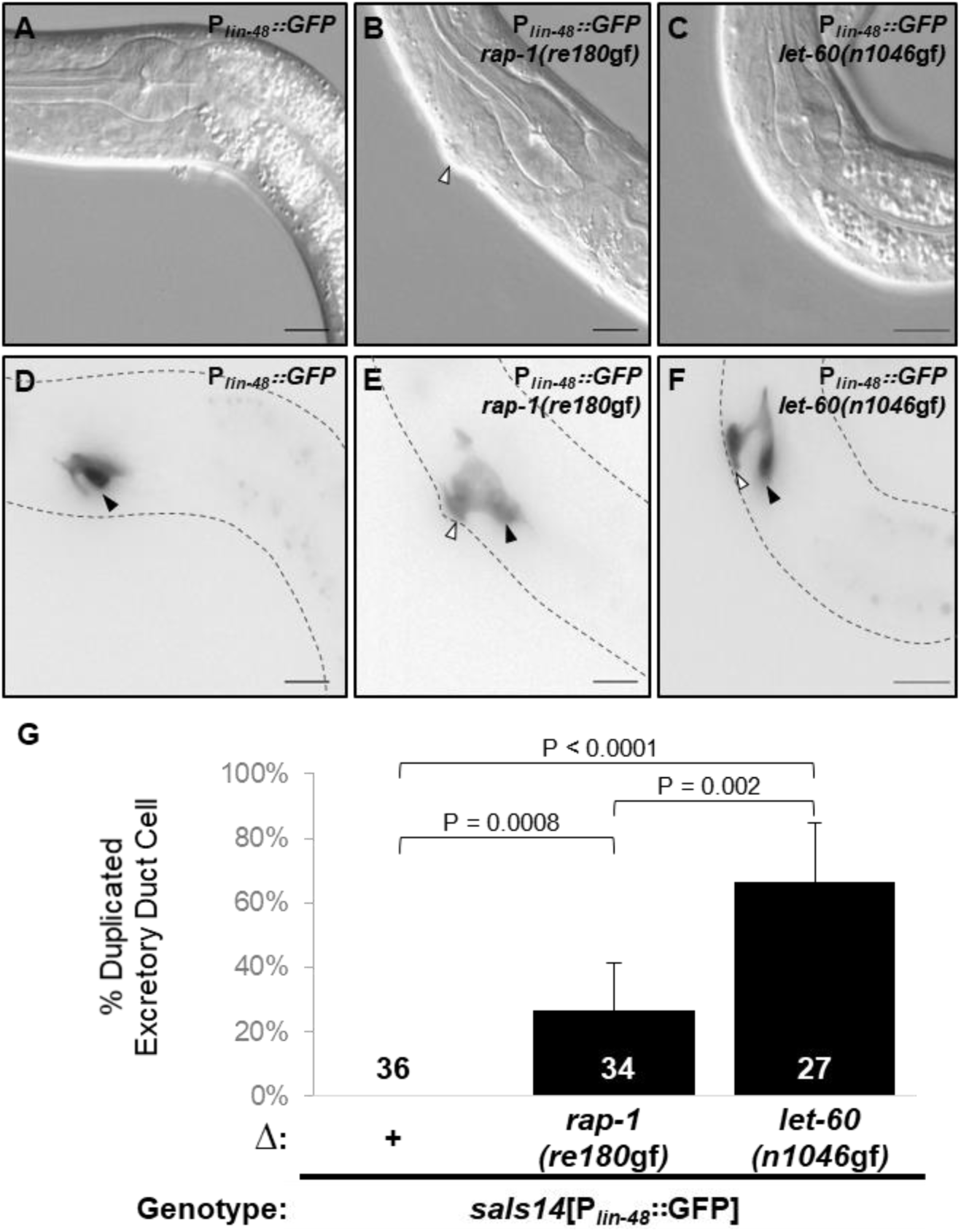
RAP-1 is sufficient to duplicate excretory duct cells. Representative DIC **(A, B,C)** and inverted epifluorescence **(D,E,F)** micrographs of excretory duct cell fate marker *saIs14*[P*lin-48::GFP*] at the L1 stage. Wild-type animals **(A,D)** have a single duct, but both *rap-1(re180*gf) **(B, E)** and *let-60(n1046*gf) **(C, F)** are sufficient to duplicate the excretory duct. Scale bar = 10 µm. **G)** Counting animals with two cells positive for P_*lin-48*_*::GFP* expression indicated that both activated *rap-1* and *let-60* resulted in significantly increased duct cell duplications compared to the wild type. Numbers in columns = N. Error bars = 95% confidence interval based on sample size, P values were calculated by Fishers Exact -test.

### PXF-1/RapGEF is required for maximal induction of 1˚ cells

The Ras superfamily of small GTPases are tightly regulated by GEFs, which promote GTP loading and hence active conformation, and by GAPs, which promote GTP hydrolysis to GDP and hence inactive conformation (Wennerberg et al. 2005). While certain mammalian GAP and GEF families have overlapping specificity for both Ras and Rap1, each also maintains unique regulators. Our results (Fig. 3D, above) suggest that GAP-1/GAP1 functions to repress RAP-1 as well as the published LET-60/Ras (Hajnal et al. 1997; Stetak et al. 2008).

Vertebrate C3G, CA-GEF, EPAC and PDZGEF have been identified as GEFs specific for Raps (Guo et al. 2016; Raaijmakers and Bos 2009). The *C. elegans* ortholog of PDZ-GEF, PXF-1, is essential for development. Loss of *pxf-1* function results in hypodermal defects, inability to molt, and variable onset larval lethality. Consistent with redundant roles for RAP-1 and RAP-2 in molting, loss of either individually results in live, superficially wild-type animals. However, loss of both *rap-1* and *rap-2* phenocopies loss of *pxf-1*, suggesting that PXF-1 functions as a joint GEF for both RAP-1 and RAP-2, as is seen in mammals (Pellis-van Berkel et al. 2005; Raaijmakers and Bos 2009).

To determine potential activators of RAP-1 in VPC patterning we assayed putative RapGEFs, *pxf-1* and *Y34B4A.4*, that are expressed in tissues other than neurons (Wormbase release WS262). Due to the essential role of PXF-1 in development, we again utilized VPC-specific RNAi in a sensitized background (*let-23(sa62*gf); *mfIs70[P*_*lin-31*_*::rde-1(+)*, *P*_*myo-2*_*::gfp]*; *rde-1(ne219)*), in which the RNAi defective phenotype of the *rde-1* mutation is rescued by VPC-specific expression of wild-type RDE-1 (Barkoulas et al. 2013). VPC-specific *pxf-1(RNAi)* (Fig. 7F) but not *Y34B4A.4(RNAi)* (Fig. S7A) resulted in decreased ectopic 1˚ induction. These findings suggest that the PXF-1 RapGEF promotes 1˚ fate, consistent with it functioning as a GEF that activates RAP-1 in VPCs.

**Figure 7.**
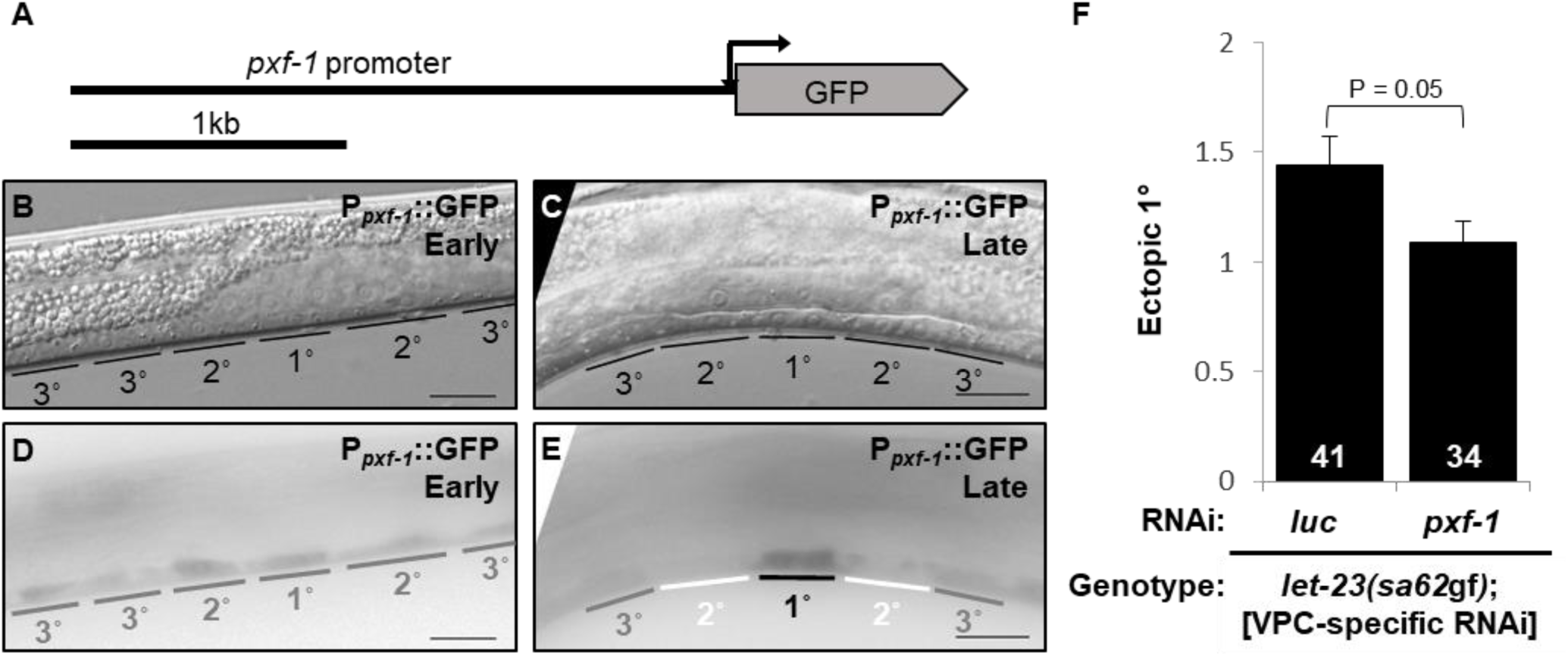
PXF-1/RapGEF contributes to 1˚ VPC induction. **A)** Diagram of the P_*pxf-1*_::GFP reporter construct. Representative DIC micrographs of L3 larvae at the **B)** early 1-cell stage, prior to EGF signaling, and the **C)** 2-cell stage, after EGF signaling. **D, E)** Representative inverted epifluorescence micrographs of the animals shown in A, B), showing GFP expression from a *pxf-1* transcriptional reporter GFP fusion transgene (*bjIs40*[P_*pxf-1*_::GFP]). **D)** GFP was observed throughout the VPCs (gray labels) prior to EGF induction at the early 1-cell stage. **E)** At the 2-cell stage, GFP increased in presumptive 1˚ (black labels) and decreased in presumptive 2˚ cells (white labels); expression in presumptive 3˚s did not change, gray labels. (scale bar = 10 µm). **F)** VPC-specific RNAi knockdown of *pxf-1* in the *let-23(sa62*gf) background decreased ectopic 1˚ induction compared to Luciferase (luc) RNAi control (*sa62*gf was cis-marked with *unc-4(e120)*). (VPC-specific RNAi = *rde-1(ne219)*; *mfIs70*[P_*lin-31*_::*rde-1*(+), P_*myo-2*_::*gfp*].). The Y-axis indicates number of ectopic 1˚s. All data are representative non-pooled assays. Error bars = S.E.M., white numbers = N. P values were calculated by T-test.

### Expression from the pxf-1 promoter is dynamically restricted to 1˚ cells

While both RAP-1 and LET-60 are sufficient to induce transformation of ectopic 1˚ cells, our results suggest such they differ both in strength and spatial activity. Unlike constitutively activated LET-60/Ras, which transforms VPCs that normally become 3˚ into 1˚ cells, constitutively activated RAP-1 mostly transforms presumptive 2˚ cells (P5.p and P7.p) to become 1˚ (Fig. 5D, Fig S6E). This observation suggested that the activity of RAP-1 is more spatially restricted within in the VPCs than is the activity LET-60, consistent with the modulatory role of RAP-1 compared to the central role of LET-60. Yet, like LET-60 (Dent and Han 1998), RAP-1 expression is uniform throughout VPCs (Fig. 2A-D). Consequently, we examined the expression of the putative GEF for RAP-1, PXF-1.

A transgenic reporter fusion of the *pxf-1* promoter to GFP (P_*T14G10.2*_::*GFP-I*), which includes - 2394 to +26 relative to the translational start codon of *pxf-1* exon 1, expressed GFP in the hypodermis (Pellis-van Berkel et al. 2005). We evaluated dynamic P_*T14G10.2*_::GFP-I expression in VPCs before and after onset of LIN-3/EGF induction using the first cell division (Pn.p to Pn.px) as a conservative temporal indicator of induction. Prior to LIN-3/EGF induction, we observed uniform GFP expression in all VPCs (Fig. 7B, C). After induction, we observed GFP expression to be increased in P6.p, the presumptive 1˚ cell, and absent in P5.p and P7.p, the presumptive 2˚ cells (Fig. 7D, E). We therefore hypothesize that *pxf-1* expression is dynamically altered in response to the onset of LIN-3/EGF expression. We propose that this regulatory mechanism serves to restrict RAP-1 activation to the presumptive 1˚ cell, while abrogating RAP-1 activation in presumptive 2˚ cells (Fig. 1).

## Discussion

Our genetic results suggest that RAP-1 functions cell autonomously to promote 1˚ fate. Our results also suggest that RAP-1, in addition to LET-60, is a substrate of the bifunctional GAP-1/GAP1. Critically, the constitutively activated endogenous RAP-1 mutant is semi-dominant and is sufficient to induce 1˚ fate. And, like *let-60(*gf), *rap-1(*gf) duplicates excretory duct cells. Yet the ectopic 1˚ induction and excretory duct cell duplication phenotypes conferred by *rap-1(*gf) are substantially weaker than those of *let-60(*gf). These observations indicate that, as expected, LET-60 plays a central role while RAP-1 plays a modulatory role. (It is worth noting that the G12V mutation that we use to activate RAP-1 was never isolated for LET-60 in screens, presumably because full LET-60 activation is predicted to confer both lethality (Schutzman et al. 2001; Pellis-van Berkel et al. 2005) and sterility (Eisenmann and Kim 1997). G13 mutations are predicted to be weaker in their activation of GTPases (Reiner and Lundquist 2016).

We also found that PXF-1/PDZ-GEF, a Rap1GEF in mammalian cells and in *C. elegans*, is required to maximally promote 1˚ fate. This function, like that of RAP-1, is also cell autonomous and suggests that PXF-1 functions as the GEF that triggers RAP-1 activation in VPCs. Activation upstream of PXF-1 remains unknown. Interestingly, like many other signaling genes in the VPC signaling network (see Introduction), expression from a *pxf-1* promoter::GFP reporter changes after initial induction. Expression prior to EGF induction is uniform throughout the VPCs, but after induction GFP signal increases in the presumptive 1˚ cell and is extinguished from presumptive 2˚ cells. Accordingly, we hypothesize that as a consequence of transcriptional reprogramming of *pxf-1* expression, the potential for RAP-1 activation is increased in presumptive 1˚ cells and decreased in presumptive 2˚ cells (Fig. 1).

RAP-1 may be activated by EGF in the same cells as LET-60, *i.e*. those VPCs closest to the EGF source, the AC. Yet we propose that the functions of LET-60 and RAP-1 are qualitatively different. The quantitatively weaker *rap-1(re180*gf) confers mostly 2˚-to-1˚ transformations (this study), while the stronger *let-60(n1046*gf) confers mostly 3˚-to-1˚ transformations: 2˚s are rarely transformed or express markers of ERK activation in the *let-60(n1046gf)* background (Beitel et al. 1990; Han et al. 1990; Berset et al. 2001; Yoo et al. 2004; Zand et al. 2011). For *let-60(n1046*gf) to transform 2˚ cells to 1˚ cells, it takes dramatic derepression of the EGFR-Ras-Raf MEK-ERK 1˚-promoting cascade, by simultaneous inactivation of 2˚-specific LIP-1/ERK phosphatase and DEP-1/EGFR phosphatase (Berset et al. 2005) or reduction of 2˚-promoting signals (Yoo et al. 2004; Yoo and Greenwald 2005; Zand et al. 2011). These differences between LET-60 and RAP-1 beyond strength of transformation may reflect different regulatory mechanisms between the two. For example, LET-60 is thought to be activated by the RasGEF SOS-1 (Chang et al. 2000; Modzelewska et al. 2007), while our data suggest that RAP-1 is activated by PXF-1. Conversely, while GAP-1 and GAP-3/RASA1 redundantly inhibit 1˚ fate induction, mammalian isoforms of both are potentially bifunctional (Kupzig et al. 2006; Raaijmakers and Bos 2009), and their relative contributions to LET-60 and RAP-1 inactivation are unclear.

RAP-1 mutant phenotypes and re-programming of expression from the *pxf-1* promoter after EGF induction are consistent with other transcriptional re-programming events in vulva signaling. We hypothesize that many levels of such regulation impose higher developmental fidelity on the system: a collective re-configuration of signaling protein expression could refine the action of signals, thus providing the transition from a ligand gradient to discrete binary outputs. In other words, re-configuration of signals could decrease the potential for developmental noise inherent in a gradient, or contradictory signals in the same cell.

Our results are also consistent with the role of Rap1 in developmental signaling as a “shadow Ras” that echoes and reinforces the main function of Ras signaling through Raf. It is difficult, however, to prove that RAP-1 signals through LIN-45/Raf. Yet our results are consistent with this model, as are the molecular characteristics of RAP-1 as a close relative of LET-60. However, an alternative model is that RAP-1 acts to reinforce other aspects of the 1˚-promoting signal. For example, RAP-1 could promote or stabilize EGFR signaling, as *Drosophila* Rap1 has been shown to regulate EGFR membrane localization and polarity through its junctional effector Canoe/afadin/AF6 (O’Keefe et al. 2009; Baril et al. 2014). Such a function, piggybacking on the strong induction of 1˚ cells by LET-60, could induce occasional ectopic 1˚ cells.

Our study provides mechanistic insights into the function of RAP-1, the *C. elegans* ortholog of mammalian Rap1 proteins that function as putative human oncogenes. We additionally establish a role for the RapGEF PXF-1 as the putative upstream GEF for RAP-1 in VPC fate patterning, and argue that transcriptional control of the *pxf-1* promoter over time provides an example of how the VPC signaling network is dynamically re-programmed during development to prevent potentially conflicting signals. We hypothesize that PXF-1-RAP-1 functions as a modulatory parallel input to reinforce and spatially restrict LET-60-LIN-45-dependent 1˚ cell induction.

## Acknowledgments

This work was supported by NIH grants R01-GM121625 and R21-HD090707 to D.J.R, ACS PF-16-083-01 post-doctoral fellowship to N.R and a Howard Hughes Postdoctoral Fellowship from the Helen Hay Whitney Foundation to D.J.D. Some strains were provided by the CGC, which is funded by NIH Office of Research Infrastructure Programs (P40 OD010440). Wormbase was used regularly. We thank the Reiner and Arur labs for helpful discussions.

---

**Figure S1.**
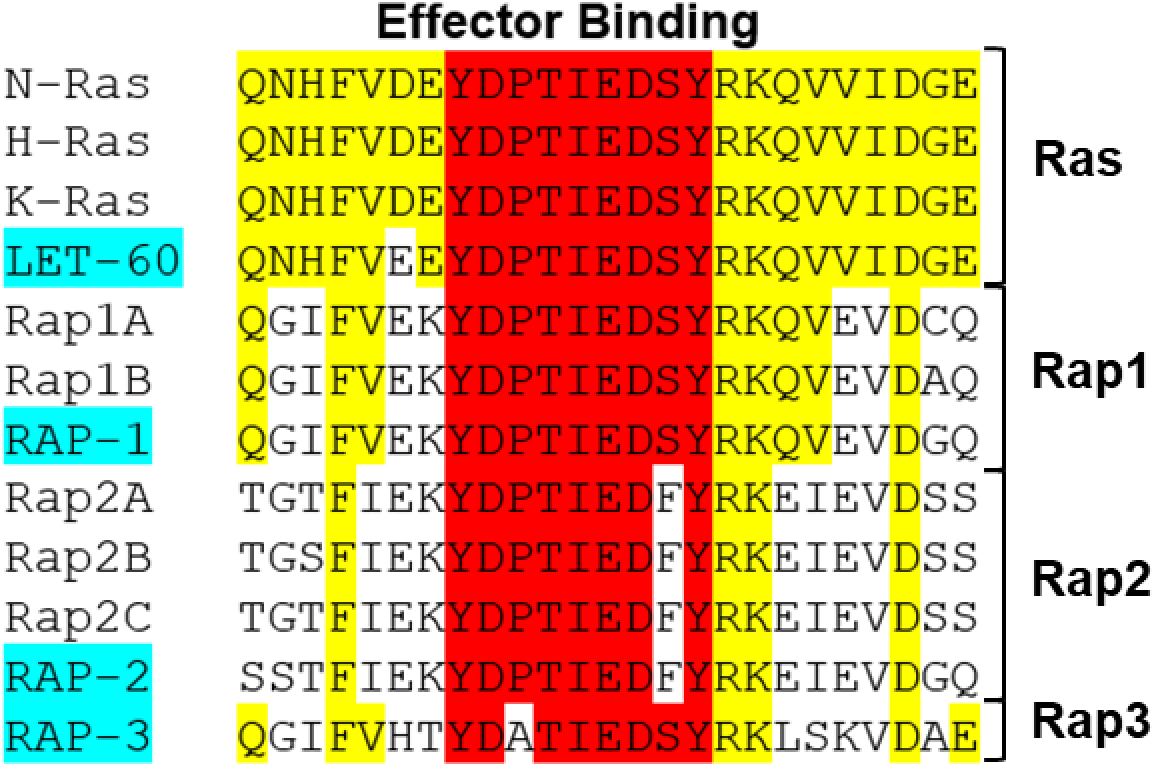
Only Rap1 shares complete Effector Binding Domain conservation with Ras. **A)** Sequence alignment of the effector binding domain of Ras, Rap1, Rap2, and Rap3 family members. The core effector domain is shaded in red and identical residues, yellow. *C. elegans* genes are shaded in blue and human genes not highlighted.

**Figure S2.**
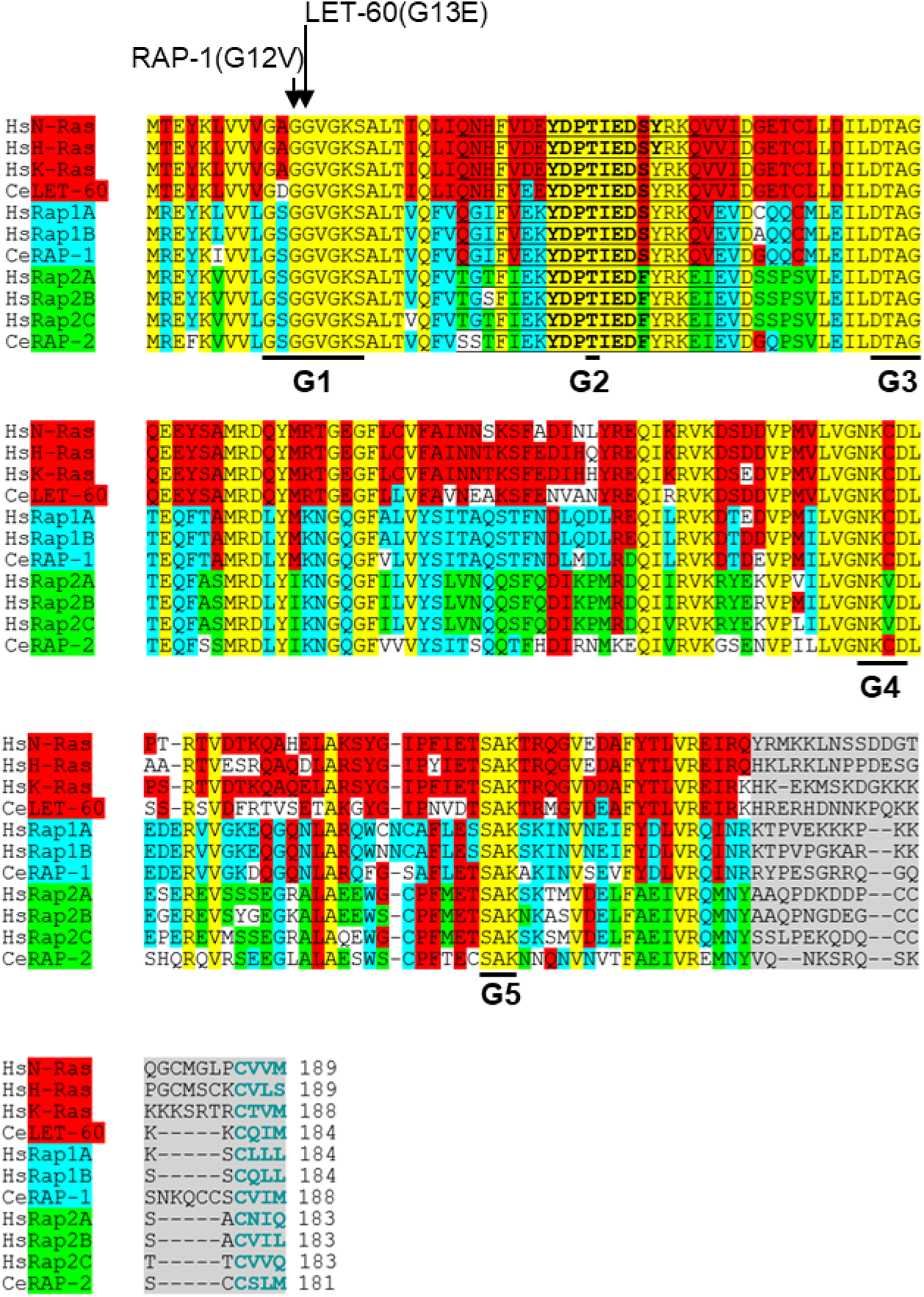
Rap1 shares greater homology with Ras than Rap2. **A)** Sequence alignment of human (Hs) and *C. elegans* (Ce) orthologs of Ras, Rap1, and Rap2. Identical residues are highlighted in yellow, with Rap1 and Rap2 conserved residues in blue and green, respectively. The effector domain of the Ras family members Is underlined with the core domain in bolded text. The C-terminal hyper-variable region is highlighted in gray and the CAAX in blue text.

**Figure S3.**
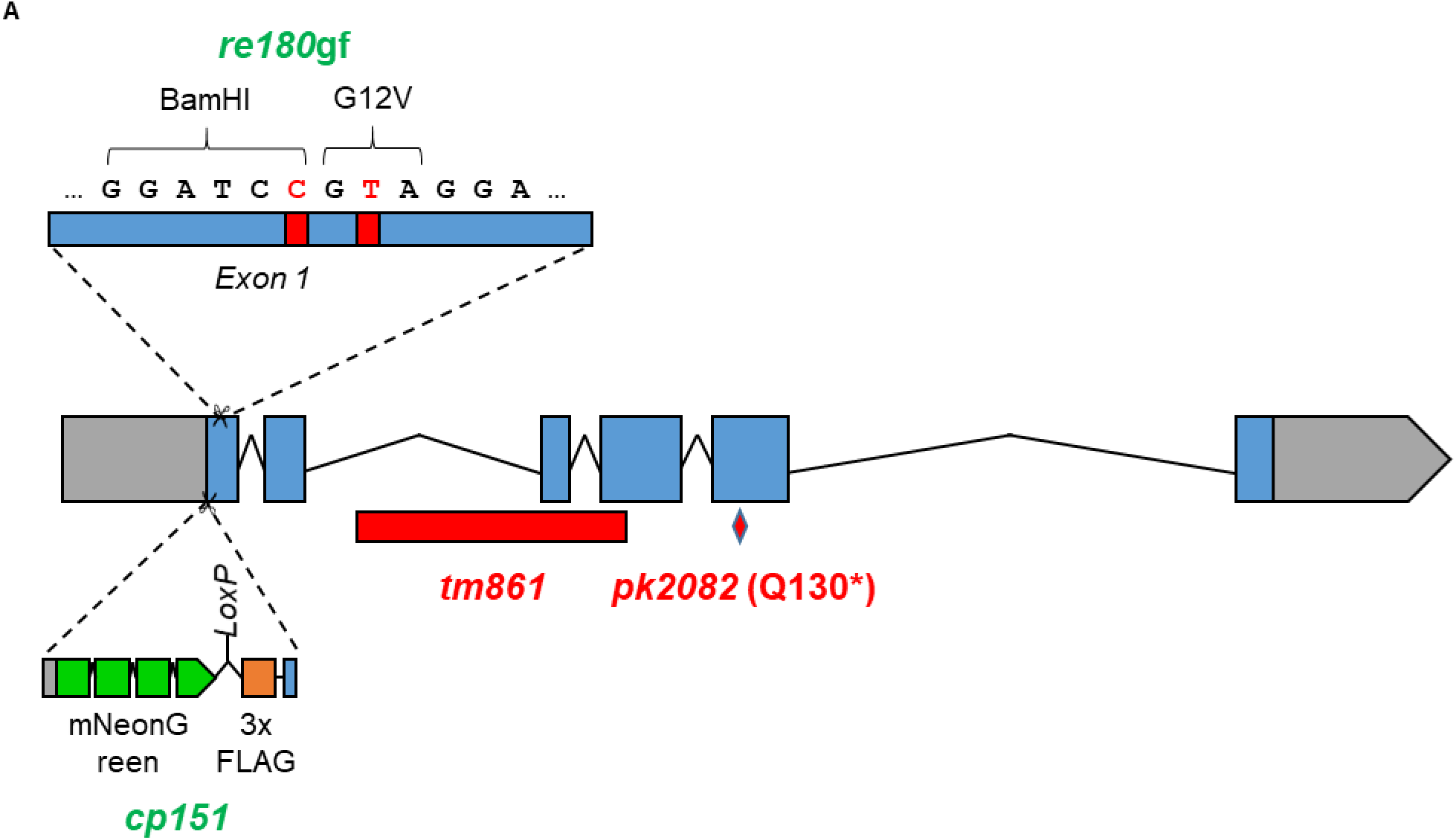
Overview of all *rap-1* alleles used. **A)** Diagram of all the *rap-1* alleles used in this study. CRISPR knock-ins are colored in green and predicted strong loss or null alleles in red.

**Figure S4.**
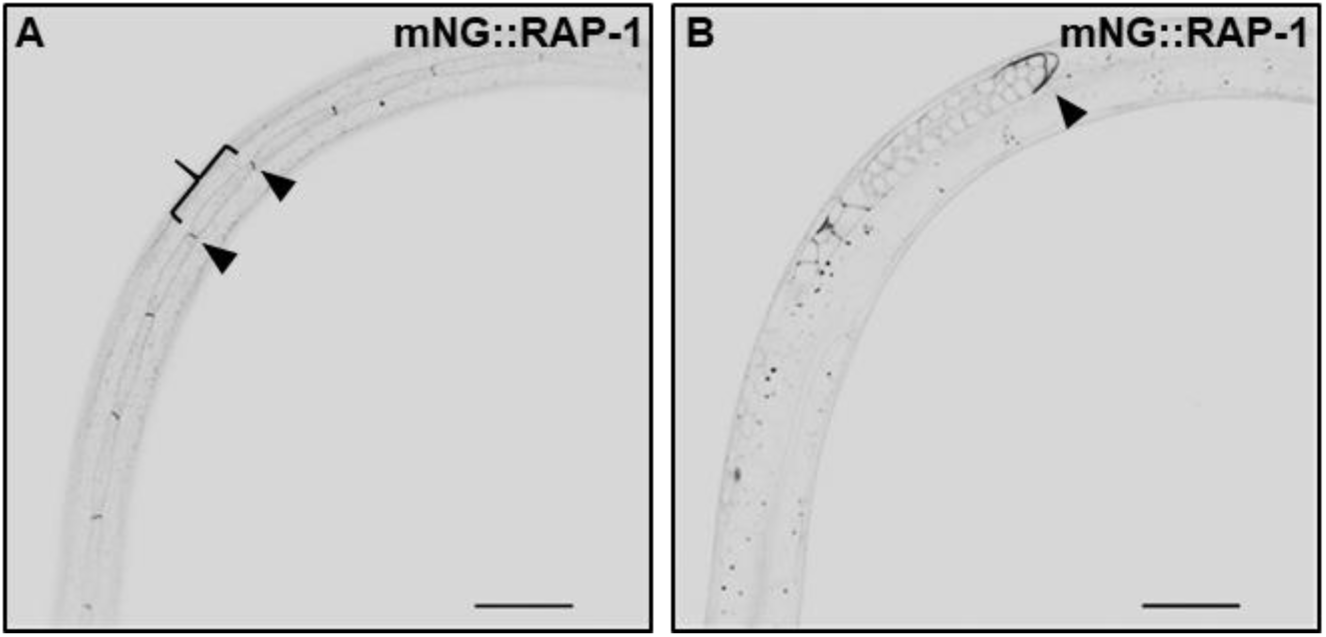
Endogenous tagged RAP-1 is ubiquitously expressed and localizes to plasma membranes and junctions. **A,B)** Representative inverted confocal fluorescence micrographs of a single *rap-1(cp151[mNG^3xFlag::rap-1])* animal at different focal planes shows expression in **(A)** hypodermal seam cells with localization at plasma membranes (bracket) and enrichment at the adherens junctions between cells (black arrowheads), **(B)** localization to plasma membranes in the germline, with increased expression in the migrating distal tip cell (black arrowhead) that invades through basement membranes. Scale bar = 20 µm.

**Figure S5.**
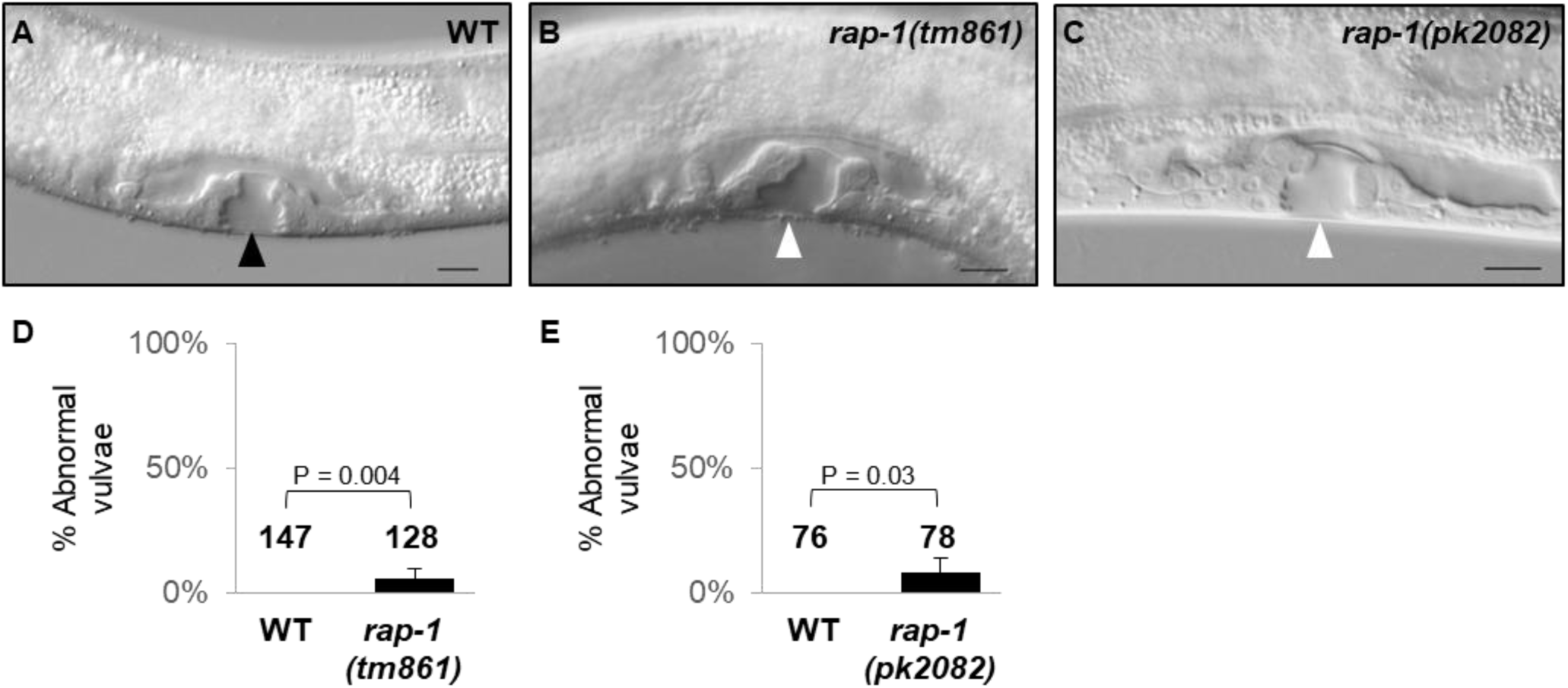
Loss of *rap-1* confers low penetrance vulval patterning defects. Representative DIC micrographs of vulvae at the late L4 stage. **A)** Wild-type vulva (black triangle). **B)** *rap-1(tm861)* and **C)** *rap-1(pk2082)* abnormal vulvae (white triangle). Scale bar = 10 µm. **D, E)** Quantification of mutant abnormal vulvae in wildtype vs. **(D)** *rap-1(tm861)* and **(E)** *rap-1(pk2082)*, with percent abnormal vulvae on the Y axis. All data are representative non-pooled assays with the mean shown. Numbers in columns = N. Error bars = 95% confidence interval based on sample size, P values were calculated by Fishers Exact-test.

**Figure S6.**
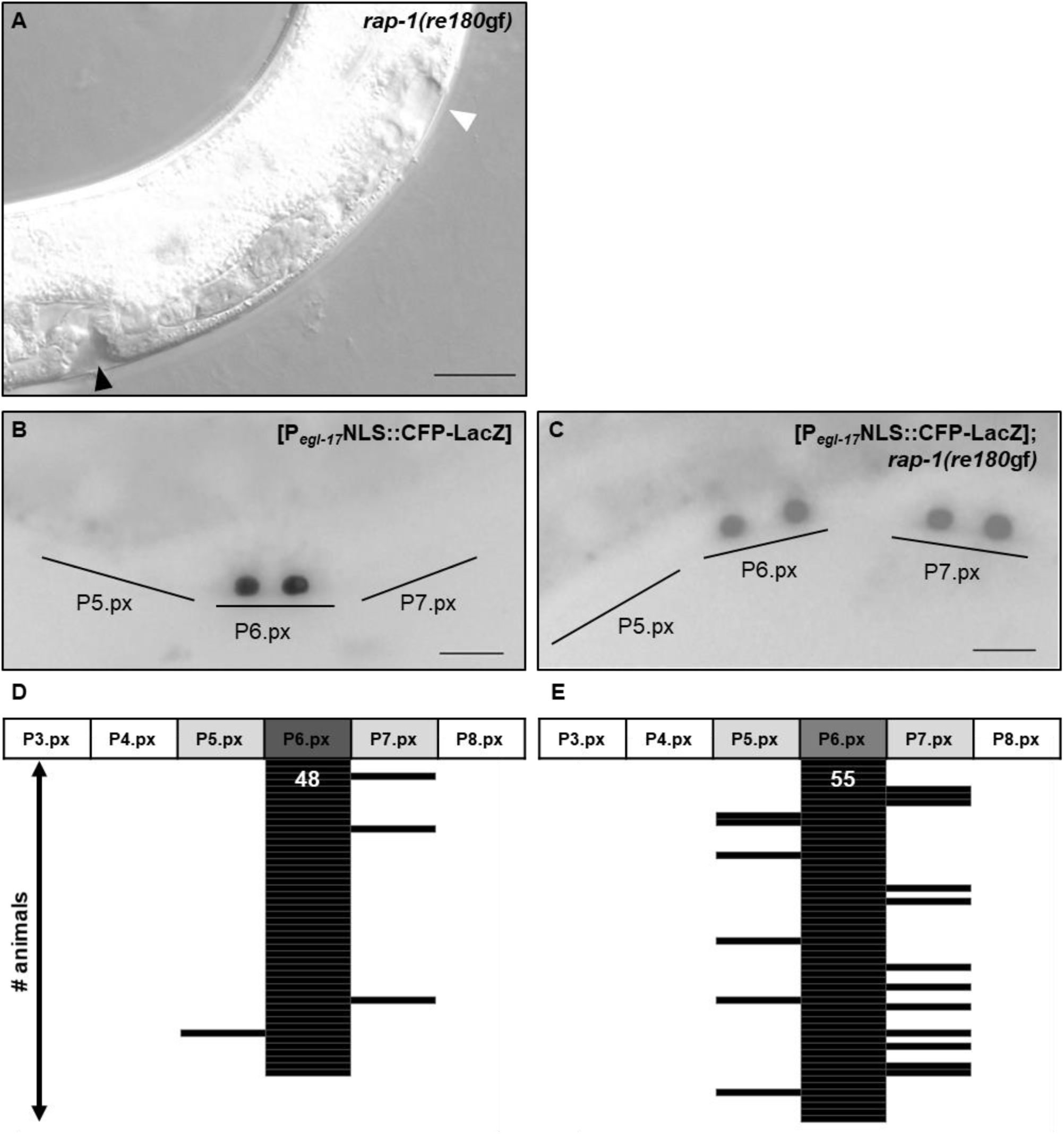
*rap-1(re180*gf) is sufficient to induce ectopic 1° VPCs. **A)** Representative DIC micrographs at the late L4 stage shows the wild-type vulva (black triangle) and induction of ectopic 1°s (white triangle) in *rap-1(re180*gf) animals (scale bar = 100 pixels). **B, C)** Representative epifluorescence micrographs in *arIs92* [P*egl-17*NLS::CFP-LacZ] animals at the 2-cell (Pn.px) stage show the inappropriate expression of 1° signaling reporter [P_*egl-17*_NLS::CFP-LacZ] in **(C)** *rap-1(re180*gf) but not **(B)** wild type (scale bar = 10 µm). **D,E)** Schematic representation of the expression of the *arIs92*[P_*egl-17*_NLS::CFP-LacZ] signaling reporter across the all six VPCs for both **(D)** control and **(E)** *rap-1(re180*gf). The addition of *rap-1(re180gf)* resulted in a significant increase of expression outside of P6.px (P = 0.003) P values were calculated by Fisher’s Exact Test. N = white number, with each line representing an animal.

**Figure S7.**
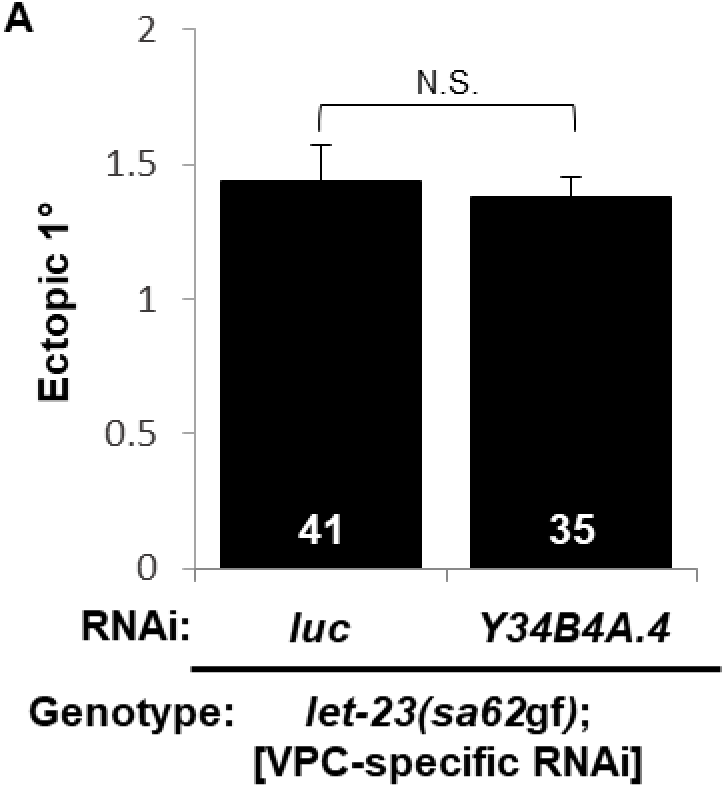
Y34B4A.4/Rap1GEF does not contribute to 1˚ VPC induction. **A)** VPC-specific RNAi knockdown of *Y34B4A.4* (*C. elegans* ortholog of Rap1GEF) in the *let-23(sa62*gf) background had no effect compared to Luciferase (luc) RNAi control (sa62 was cis-marked with *unc-4(e120)*). (VPC-specific RNAi = *rde-1(ne219)*; *mfIs70*[P_*lin-31*_::*rde-1*(+), P_*myo-2*_*::gfp*].) The Y-axis indicates number of ectopic 1˚s. All data are representative non-pooled assays. Error bars = S.E.M., white numbers = N. P values were calculated by T-test.

**Supplementary Table 1.**
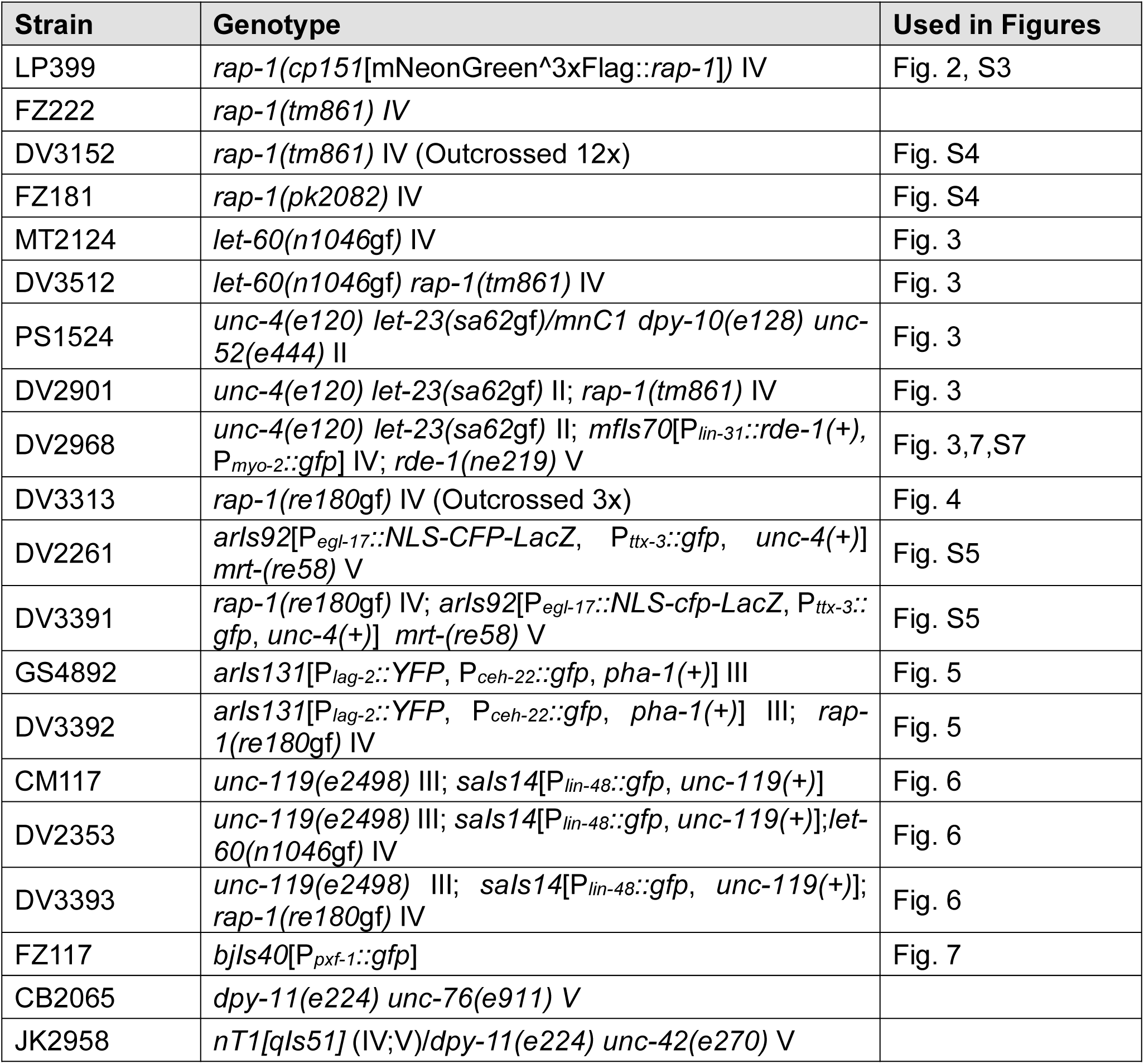
Strains

**Supplementary Table 2.**
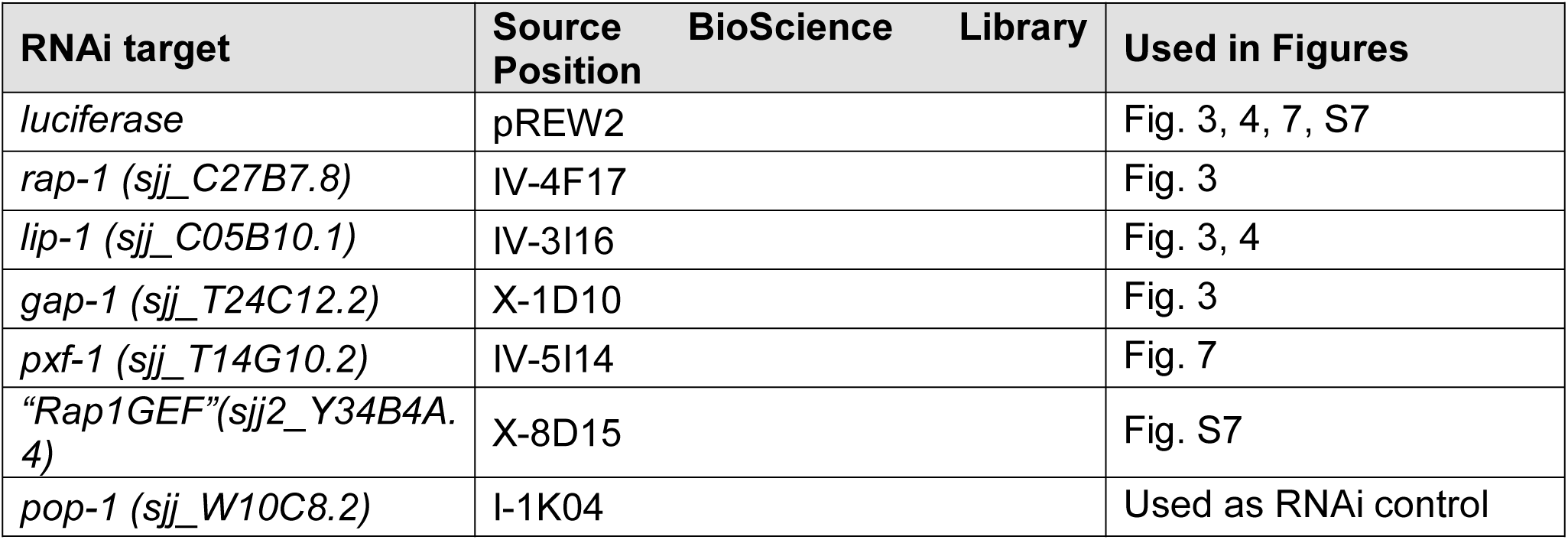
RNAi

**Supplementary Table 3.**
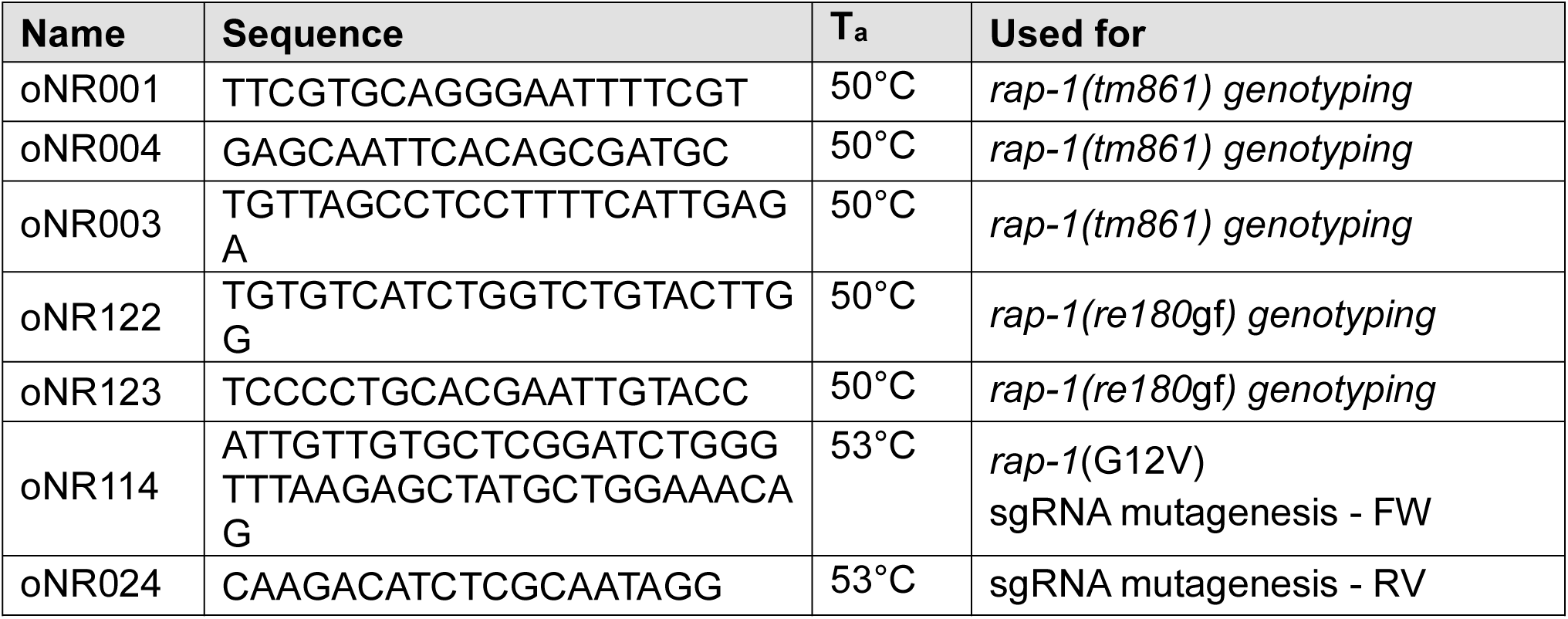
Primers

**Supplementary Table 4.**
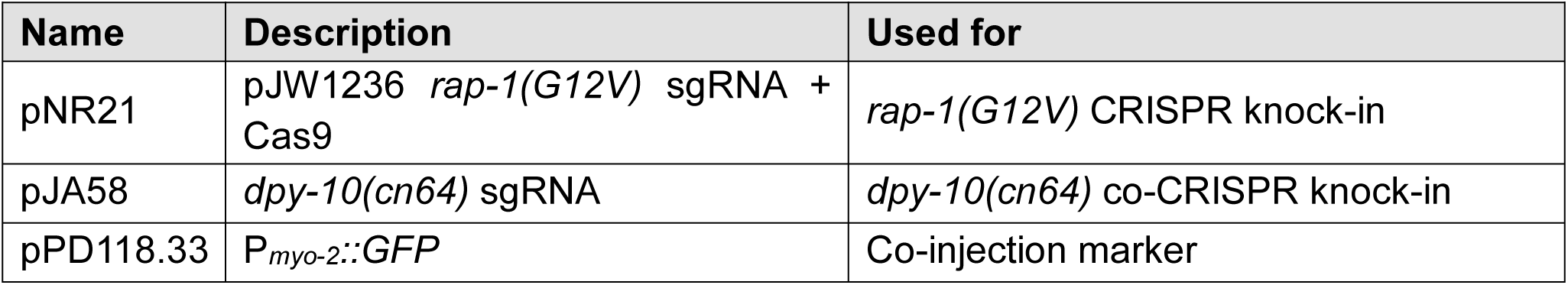
Plasmids

**Supplementary Table 5.**
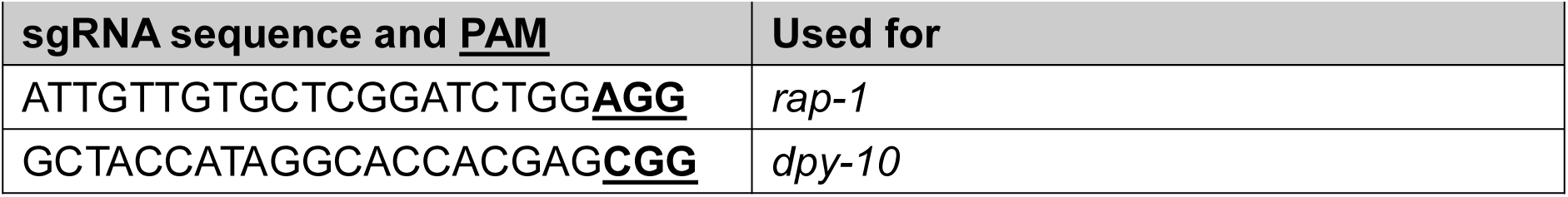
sgRNA sequences and PAMs

**Supplementary Table 6.**
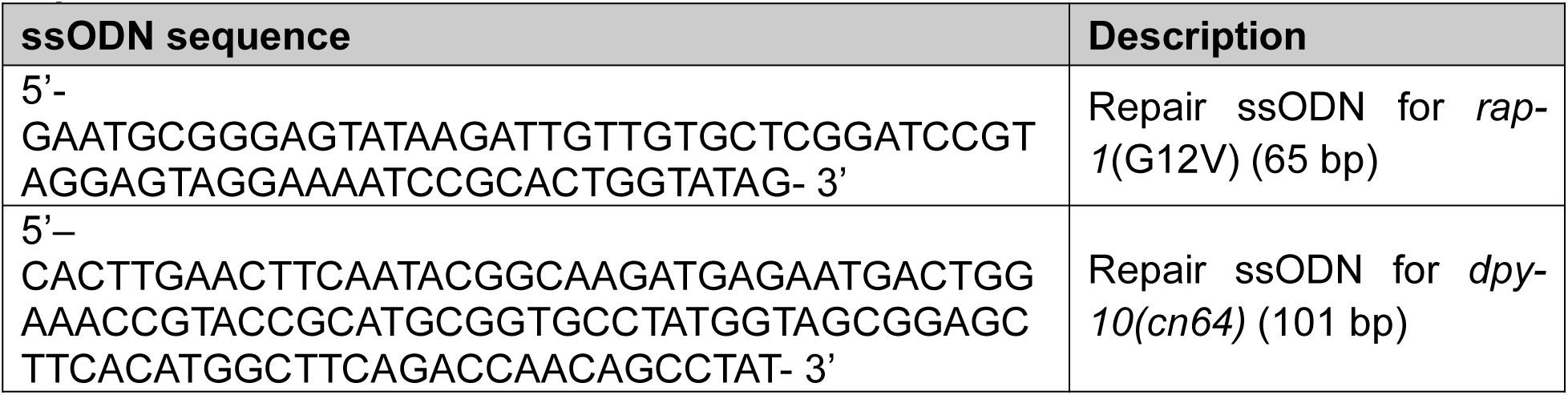
Repair ssODN

